# Occurrence-based diversity estimation reveals macroecological and conservation knowledge gaps for global woody plants

**DOI:** 10.1101/2023.04.09.536180

**Authors:** Buntarou Kusumoto, Anne Chao, Wolf L. Eiserhardt, Jens-Christian Svenning, Takayuki Shiono, Yasuhiro Kubota

## Abstract

Incomplete sampling of species’ geographic distributions has challenged biogeographers for many years to precisely quantify global-scale biodiversity patterns. After correcting for the spatial inequality of sample completeness, we generated a global species diversity map for woody angiosperms (82,974 species, 13,959,780 occurrence records). The estimated diversity demonstrated non-linear latitudinal and longitudinal patterns that were potentially related to region-specific biogeographic factors including current climate, paleoclimate, and topographical factors, while energy availability was the most important predictor at a global level. We identified the areas with potentially high species richness and rarity, but poorly explored, unprotected, and threatened by deforestation: they are distributed mostly at low latitudes across central South America, central Africa, subtropical China, and Indomalayan islands. These priority areas for botanical exploration would help to efficiently fill spatial knowledge gaps for better describing the status of biodiversity and improve the effectiveness of the protected area network for global woody plant conservation.

**Teaser:** Bias-corrected diversity map based on occurrence records sheds new light on global macroecology and conservation of woody angiosperms.

## Introduction

The accumulation of species occurrence data is a fundamental basis for biodiversity science, providing rising opportunities towards addressing major challenges in ecology and conservation (*1, 2*). Occurrence records have been widely used to model species distribution (*3*) and to estimate diversity at given localities (*4*). However, occurrence records notoriously suffer from incompleteness and biases (*5*), where observed species diversity is statistically influenced by sample size (*6*). As most occurrence records stem from collections taken for purposes other than estimating diversity patterns, their coverage is usually not geographically systematic nor comprehensive, resulting in a dominance of omission errors (*7, 8*); to complicate matters, species absence is scale-dependent, and its information is usually unavailable (*9*). This so-called Wallacean shortfall in biodiversity knowledge (*10*) potentially precludes a solid understanding of geographical biodiversity patterns (*11, 12*) and implementation of spatial conservation planning (*13*).

To correctly capture species diversity patterns, knowing the geographic variation in sample completeness of species occurrence data is critical (*14*). The explicit link between sample size, completeness, and diversity enables standardization of an observed diversity using rarefaction or extrapolation based on sample completeness (*15*). This allows fair comparisons of species diversity across multiple assemblages measured at unequal sample completeness without necessarily knowing their true diversity (*14*). Notably, latitudinal and longitudinal diversity gradients have recently been revisited in this manner, especially in marine ecosystems (*16, 17*) and revealed unexpected diversity patterns (e.g. bi- or multi-modality). Thus, diversity estimation theory sheds light on the generality of macroecological patterns that often suffer from serious sampling bias (*18*).

To achieve the global goals and milestones to counteract the current biodiversity crisis (e.g., the post-2020 Biodiversity Framework; *19*), the spatial allocation of conservation resources (e.g., land areas) is a key issue. For effective avoidance or mitigation of negative human impacts, spatial planning based on reliable information of biodiversity distribution is essential (*20*). However, spatial planning analyses implicitly assume that biodiversity is described precisely/equally both inside and outside of existing conservation areas; the validity of the assumption is not examined on a global scale yet.

In this study, we focused on the species diversity of woody angiosperms. Woody angiosperms play a crucial role as ecosystem engineers, shaping most terrestrial biomes and supporting ecosystem functions and services on Earth (*21*). A recent study applied diversity estimation theory to a global occurrence record dataset, and estimated the continental-level tree species richness with correcting uneven sample completeness (*4*). However, their analysis did not include all woody angiosperms, and only estimated diversity at the level of bioregions (biomes on continents). Global patterns of woody plant diversity at finer resolutions remain to be estimated from occurrence records, and compared to previous studies using different data sources such as floristic checklists (*22, 23*) and plot surveys (*24*).

Here, we generated a global diversity map for woody angiosperms using 13,959,780 occurrence records for 82,974 species. Sample completeness and standardized species diversities using a Hill number-based approach were computed to examine bias-corrected geographical patterns of species diversity. Hill numbers (or the effective number of species) (*25*) have been increasingly used to quantify the species diversity of assemblages. In particular, we evaluated the impact of sample completeness on the description of species diversity and ecological inferences, especially of latitudinal and longitudinal diversity patterns related to spatial resolution, and identified predominant environmental drivers of species diversity at global and regional scales. We also examined the spatial congruence of the species diversity and sample completeness with the global protected area network, and the deforestation trend during 2000 to 2020. Finally, we identified spatial priority areas for allocation of future sampling effort to effectively fill knowledge gaps.

## Results and Discussion

### Observed diversity and sample completeness

Sample completeness measured by sample coverage, a concept originally developed by Alan Turing in his cryptographic analysis during World War II, greatly varied globally for the occurrence records of woody angiosperms (Fig. 1) (*5, 26, 27*). Sample coverage tended to be high in temperate regions, including North America, Europe, Japan, Australia, and New Zealand. Such geographical inequality of sampling effort distorts the description, interpretation, and prediction of biodiversity patterns (*27*–*29*), because the observed diversity patterns reflect both multiple gradients of true diversity and the spatial bias of sampling efforts (*30*). Indeed, the observed number of species showed a strong spatial congruence with the total number of occurrences (Fig. S1). Expectedly, sample coverage was lower and more variable at the finest spatial resolution (100 km × 100 km) than at coarse resolution (∼ 800 km × 800 km) (Fig. S2). Such positive scale-dependency of sample completeness has been reported previously in a regional-scale study of plants (*31*) and a global-scale study of stony corals (*17*).

**Fig. 1.**
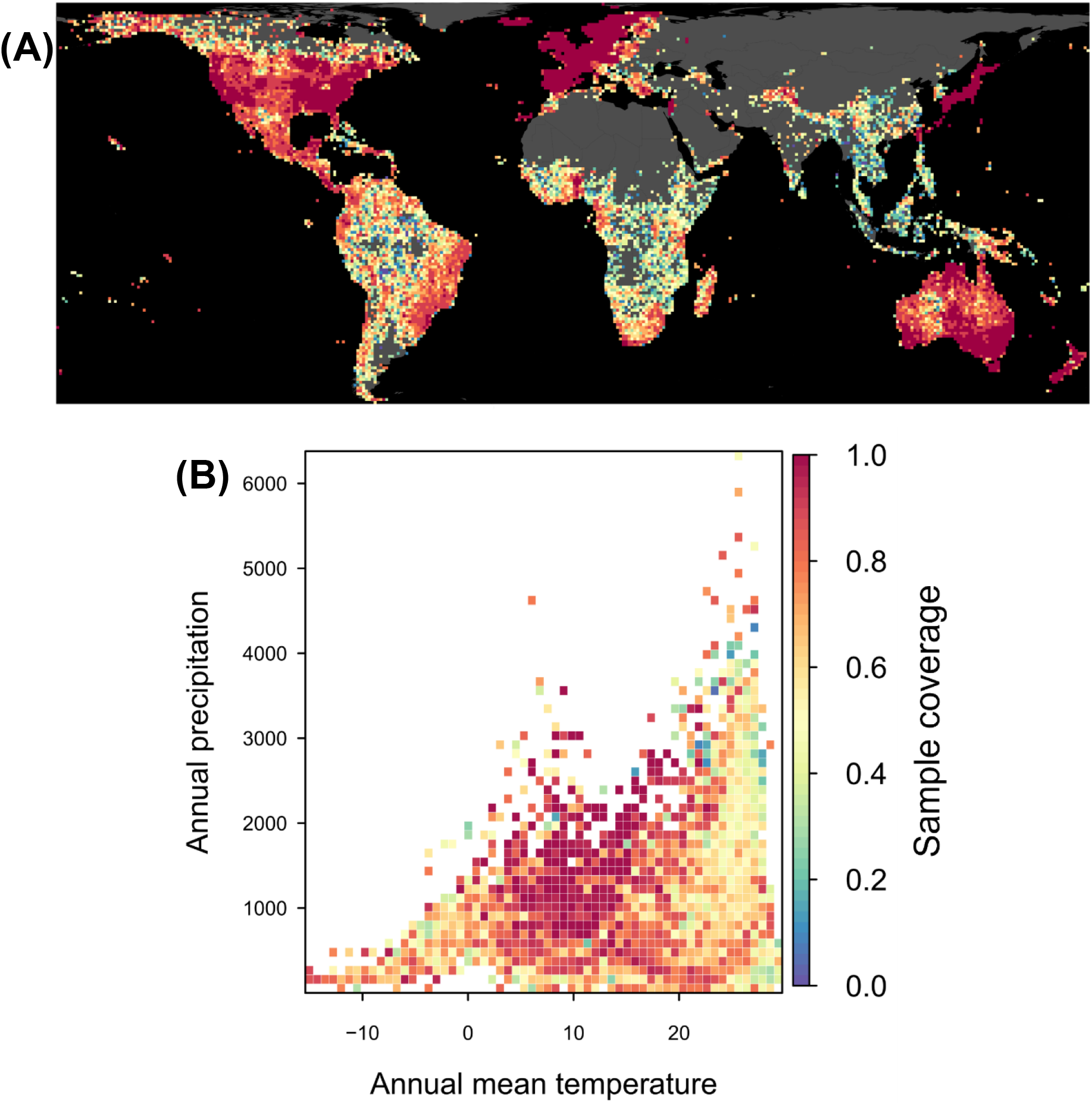
Sample completeness (sample coverage) of species occurrence records of woody angiosperms at global scale: **(A)** geographical map at the 100 km × 100 km equal-area grids and **(B)** the distribution in a climograph plot.

The sample coverage-based standardization of species diversity successfully mitigated the effect of uneven sample completeness (*14*), improving the description of relative geographical diversity patterns (Figs. 2–4, S3–S9). Notably, the improvement is quantitatively evident from the increase in the predictive performance of macroecological models to explain species richness patterns after standardization (Figs. 5, S10 and S11).

**Fig. 2.**
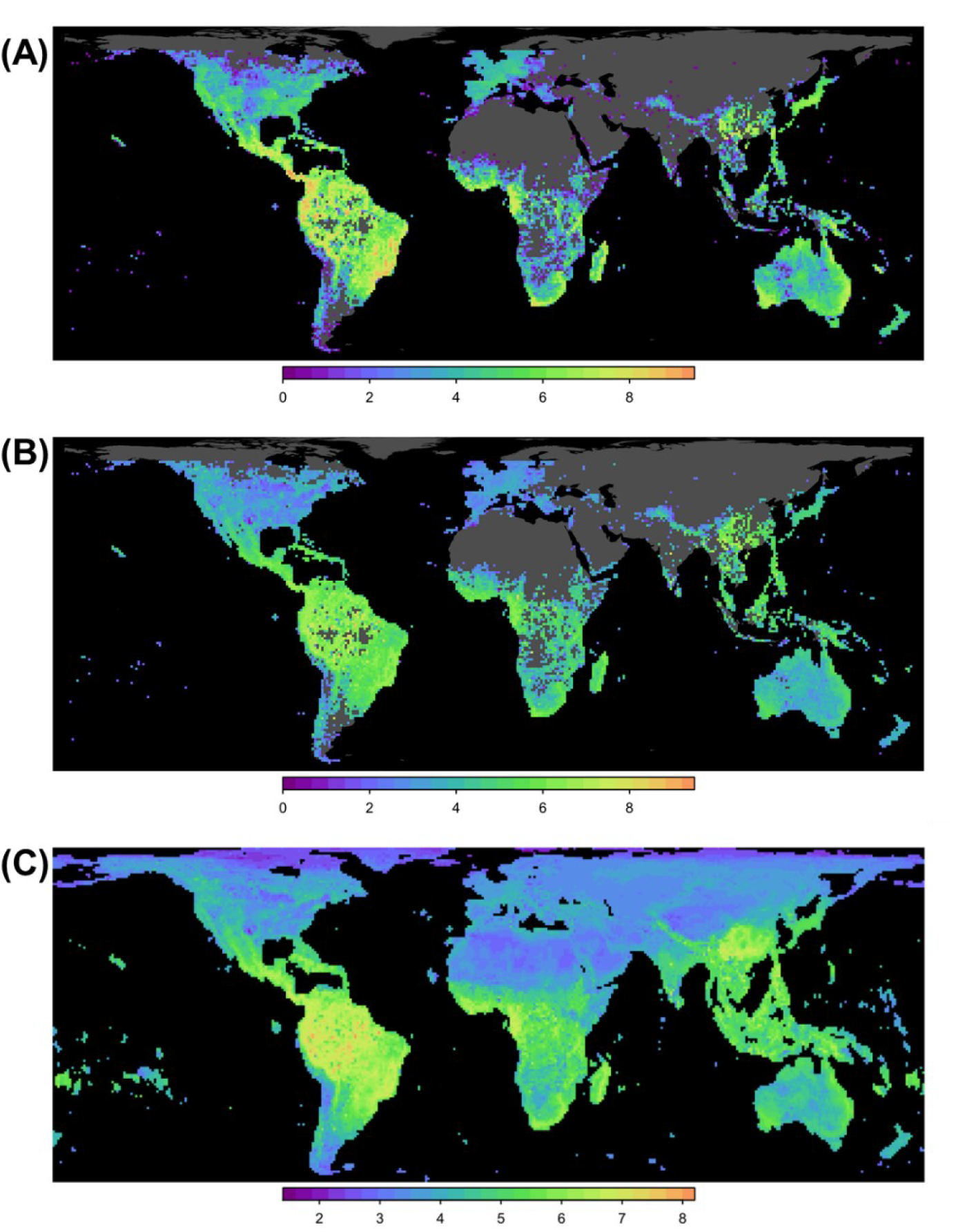
Geographical distribution of species richness (Hill number *q* = 0) for 100 km × 100 km grid cells: **(A)** observed species richness, **(B)** sample coverage-based standardised species richness (sample coverage = 0.82); **(C)** spatial projection of Random Forest model for standardized richness. The values are log-scaled. Patterns for diversity at higher orders (*q* = 1, 2) are given in Fig. S18.

**Fig. 3.**
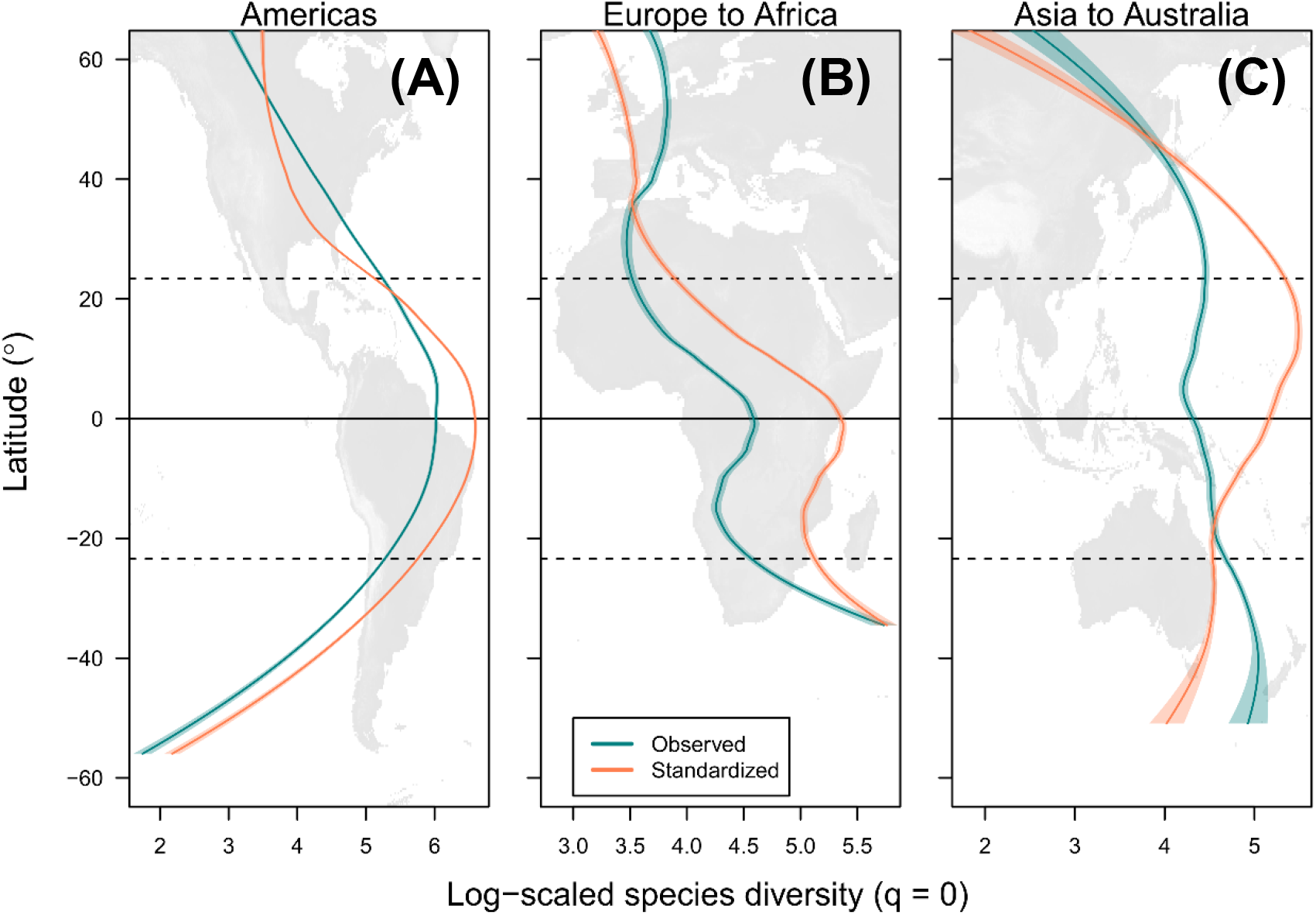
Latitudinal pattern of species richness in three longitudinal zones. The globe was subdivided into **(A)** North-to-South Americas, **(B)** Europe-to-Africa, and **(C)** Asia-to-Australia: the observed species richness and the standardized species richness (species diversity at the order *q* = 0) based on sample coverage (0.82). Loess (locally estimated scatterplot smoothing) curve (scaling parameter alpha = 0.6) is shown (green and red lines). Thick horizontal line indicates the equator, and dashed lines represents the Tropics of Capricorn and Cancer. Patterns for diversity at higher orders (*q* = 1, 2) are given in Fig. S19.

**Fig. 4.**
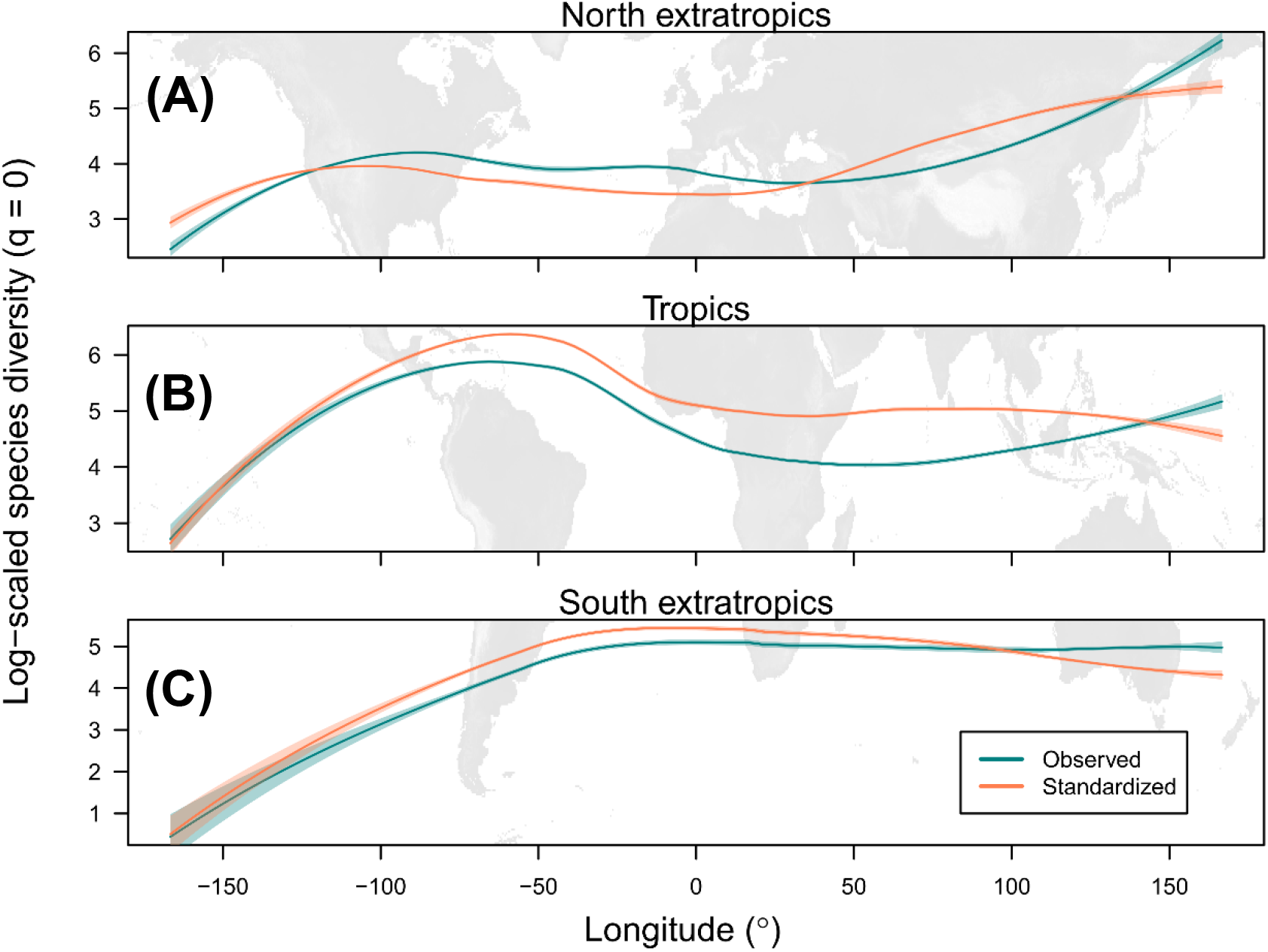
Longitudinal pattern of species richness in three latitudinal zones. The globe was subdivided into **(A)** northern extratropics, **(B)** Tropics, and **(C)** southern extratropics: the observed species richness and the standardized species richness (species diversity at the order *q* = 0) based on sample coverage (0.82). The diversity values are log-scaled. Loess (locally estimated scatterplot smoothing) curve (scaling parameter alpha = 0.6) is shown (green and red lines). Patterns for diversity at higher orders (*q* = 1, 2) are given in Fig. S20.

Specifically, the best-performing Random Forest model explained ∼75% of the total variance of the sample coverage-based standardized species richness pattern, while for the observed species richness the model explained only 63% of the total variance (Figs. 5a and Fig. S10). This is comparable to previously reported values (70%–85%) in studies of macroscale plant diversity (*23, 24, 32*–*34*). These improved estimates of species richness showed deviations from the geographical patterns of observed species richness (Figs. 2–4, S12) especially for the latitudinal gradients in the Asia-to-Australia region (around the Tropic of Cancer) and in the Europe-to-Africa region (around equator), likely reflecting that the observed species diversity is severely affected by undersampling. Furthermore, the deviations in geographic patterns between observed and standardized diversities were scale-dependent and tended to be greater at finer spatial resolutions (Figs. S13–S17).

**Fig. 5.**
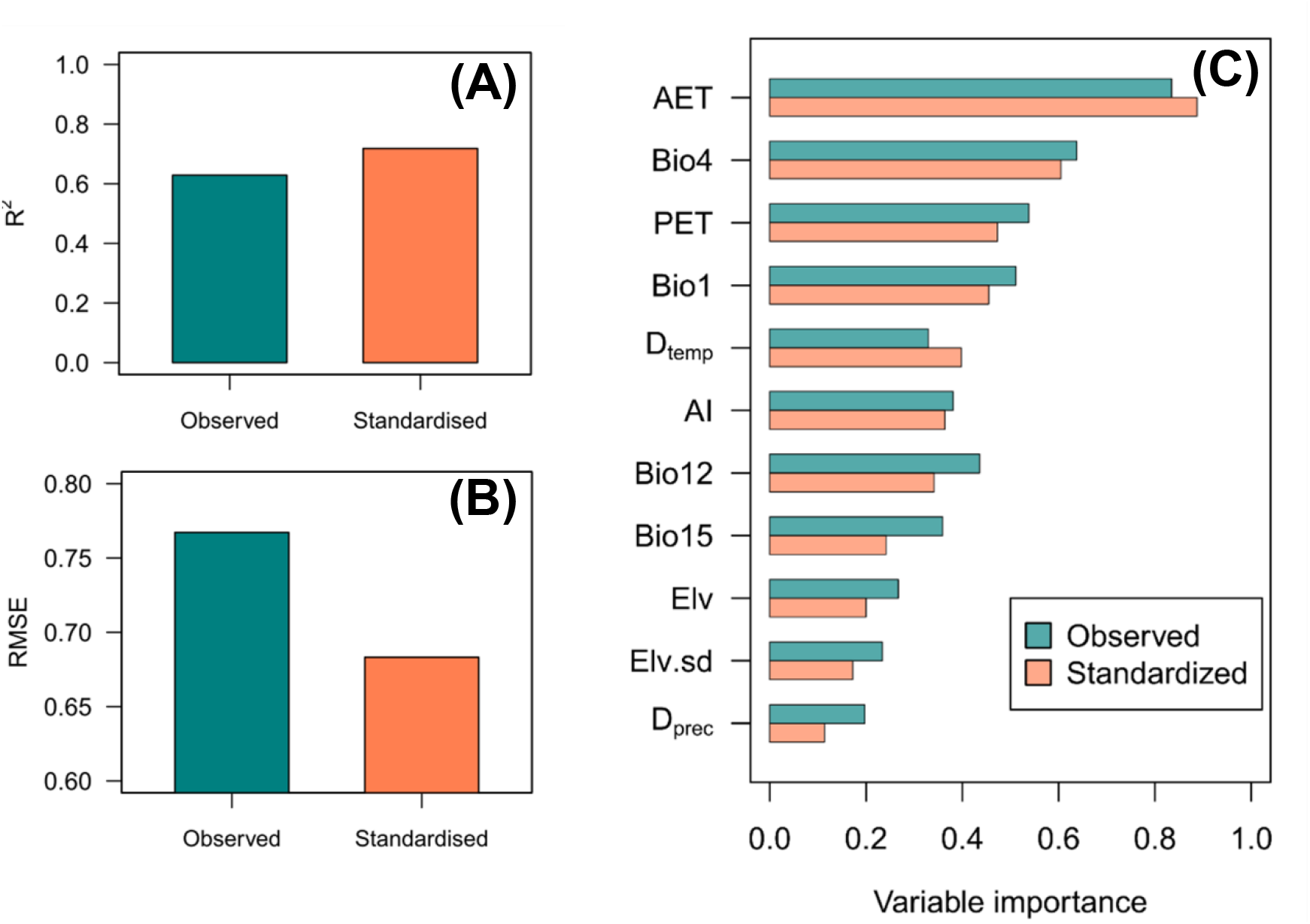
Outputs of the Random Forest model explaining the observed and the sample coverage-based standardized species richness (sample coverage = 0.82): **(A)** explanatory power (*R*^2^), **(B)** root mean square error of prediction (RMSE), and **(C)** the relative importance of environmental variables. The environmental explanatory variables are mean annual temperature (Bio1), temperature seasonality (Bio4), annual precipitation (Bio12), precipitation seasonality (Bio15), actual evapotranspiration (AET), potential evapotranspiration (PET), aridity index (AI), average elevation (Elv), standard deviation of elevation (Elv.sd), and differences in temperature (D_temp_) and precipitation (D_prec_) between the Last Glacier Maxima and the present. Results for diversity at higher orders (*q* = 1, 2) are given in Figs. S21 and S22.

After applying standardization, the different orders of diversities (*q* = 0: species richness, *q* = 1: Shannon diversity, *q* = 2: Simpson diversity) showed similar geographical pattern (Figs S18–20). To our knowledge, this is the first global scale map for woody angiosperm (or vascular plants in general) diversity of higher diversity orders. Because the rarefaction/extrapolation estimators for higher orders (*q* = 1 and 2) are nearly unbiased and valid for a wide range of prediction (*15*), the consistent results among different orders of diversities suggest the standardization of species diversity improved descriptions of geographical diversity patterns.

### Geographical patterns of estimated diversity

The standardized (bias-corrected) species richness showed global latitudinal trends with regional diversity hotspots (areas characterized by high species diversity) mainly scattered in the tropics (Fig. 2). The highest diversity was identified in central South America followed by western tropical Africa and Indomalayan-Australasian region including subtropical China; these are generally in line with the estimation for tree species at continental and biome levels by Gatti et al. (*4*), while our analysis also captured finer-scale variation in species richness within continents, including a species diversity peak in the northern mid-latitudes of Asia.

The sample coverage-based standardization changed the shapes of the latitudinal diversity gradient in the three longitudinal zones (Americas, Europe-to-Africa, and Asia-to-Australia) (Figs. 3 and S3), indicating that the biased raw data lead to mischaracterization of the latitudinal diversity gradient (*18*). After bias correction, the latitudinal diversity gradients showed regional differences (Fig. 3). The latitudinal diversity gradient in Americas showed a typical symmetric shape with a peak at the equator and decline toward both poles, in line with the finding of meta-analyses (*35, 36*). In contrast, the latitudinal diversity gradient in the Europe-to-Africa and Asia-to-Australia regions showed complex patterns with asymmetric unimodality or bimodality. Specifically, the latitudinal diversity gradient in the Europe-to-Africa region showed a peak at the equator and decline toward higher latitude in the Northern Hemisphere and up to middle (∼20 degrees) latitude in the Southern Hemisphere; however, an extraordinary high species diversity at the Cape Floristic Region in South Africa (*37*) resulted in a distinctive bimodal diversity pattern (Fig. 3b). The latitudinal diversity gradient in the Asia-to-Australia region showed a peak at northern central latitudes (around the Tropic of Cancer). These regional anomalies of the latitudinal diversity gradients have been pointed out for all vascular plants using different data sources (*38*).

These region-specific latitudinal diversity gradients can be understood in the context of a biodiversity anomaly among regions with similar climatic conditions (*39*), e.g., the diversity depression of African tropical rain forests in comparison with other tropical biomes (*40*) and the ‘Asian bias’ in species diversity of temperate floras (*41*). Such regional diversity anomalies among continents were reflected in the longitudinal diversity gradients globally within the tropics and extratropics (Fig. 4 and S4). Uni- and bimodal longitudinal diversity gradients were identified in the three latitudinal zones (northern extratropics, tropics and southern extratropics). A tropical diversity peak in South America and relatively lower diversity in the rest of tropics (sub-Saharan Africa, South-East Asia, Australia and Oceania) was evident, as reported in a continental-level estimation (*4*) and community-based studies (*42-44*). Furthermore, temperate East Asia was identified as a diversity peak on the longitudinal diversity gradient in the northern extratropics, as argued in previous studies (*39, 43*), although the diversity pattern was scale-dependent and assumed a bimodal shape at coarse spatial resolutions (≥400 km × 400 km). We noted that the longitudinal diversity gradients were not steep (except for oceanic gaps) and were relatively stable irrespective of the sampling bias (Figs. 4 and S4) compared with the latitudinal diversity gradients.

### Environmental drivers of woody species diversity

Within the 11 environmental (mostly climatic, see Materials and Methods for details) factors considered, actual evapotranspiration (AET) was the most important factor to explain the geographical patterns of species richness at the global scale (Fig. 5c), regardless of the modelling framework and spatial resolution (Figs. S8–S9): AET was consistently positively associated with species richness (Figs. S5–S7), as reported in previous global-scale studies using different types of data (*23, 24 32, 34, 45*). The other climatic variables also were of relatively high importance. Interestingly, temperature change from the Last Glacier Maximum (LGM) was of a great importance in shaping species diversity, especially at coarser spatial resolutions (Fig. S9), suggesting that geographic variation in late-Quaternary paleoclimate instability has had a regional effect on macroscale diversity patterns in woody angiosperms through extinction and dispersal limited range dynamics (*46*–*49*). This is in line with the findings for local plant communities on a global geographical gradients by Sabatini et al. (*50*) where the geohistorical effects on species diversity were greater at coarser grain sizes. In addition, the relative importance of environmental drivers, especially for the top four variables, was stable across the tested spatial resolutions (Fig. S9). Such a grain-size-independent relationship of environmental variables with species diversity, which contrasts with the findings of Keil & Chase (*24*) covering a wider range of grain size across local communities (10^−3^ km^2^ ∼) to regional species pools (∼ 10^6^ km^2^), suggests the predominance of environmental species sorting at the regional species pool level (>10^4^ km^2^).

The latitudinal and longitudinal diversity gradients had region-specific links to different environmental variables (Figs. S23 and S24). Temperature seasonality outperformed energy variables (e.g., AET) to explain the species diversity in the Americas and the Europe-to-Africa region, but not in the Asia-to-Australia region (Fig. S24), suggestive of region-specific species sorting associated with latitudinal climatic seasonality gradients (*51*). In addition, historical temperature change from the LGM contributed substantially to shaping the latitudinal diversity gradient in Americas. This suggests the long-lasting impact of glacial disturbance and climatic disequilibrium in North America (*52, 53*): Seliger et al. (*52*) found that the potantial ranges of North American trees and shrubs are largely unfilled due to dispersal lags in response to post-glacial warming. Our result is also consistent with the finding of a strong correlation between historical climatic stability and number of restricted-range vertebrates in Meso-to South America (*54*). However, across all continents the longitudinal diversity gradients were mainly driven by energy and/or water variables (Fig. S24). Especially in the tropics, potential evapotranspiration and aridity index were of particularly high importance, indicating that low water availability depresses species diversity in particular tropical forests (*55*). In contrast to the latitudinal diversity gradients, topography and historical climatic changes from the LGM played minor roles in explaining the longitudinal diversity gradients (Fig. S24).

### Spatial priority areas for future sampling

We found that woody angiosperm diversity had been explored with similar sample completeness across inside and outside the protected areas in general, except for the Eastern Palearctic region where sample coverage was much lower in unprotected areas (Fig. S25). The areas with low sample coverage partly overlapped with high species richness and rarity areas which were distributed inside and outside the protected areas (Fig. 6 and S25a, b). In addition, the spatial prediction of the Random Forest model (see above) demonstrated that some sites with no occurrence data (where the species diversity estimation was impossible) could contain areas with high species diversity (Fig. 6). More importantly, those less or not explored sites are potentially suffering from deforestation (Fig. S25d). The threat of deforestation is likely to be more prominent in unprotected forests than in protected ones because of its absence of legal restrictions and deforestation leakage from surrounding protected areas (*56, 57*). Our recommendation would be that developments should be refrained in such data-deficit areas to avoid unexpectedly large biodiversity loss, including the extinction of scientifically undescribed species (*58*).

**Fig. 6.**
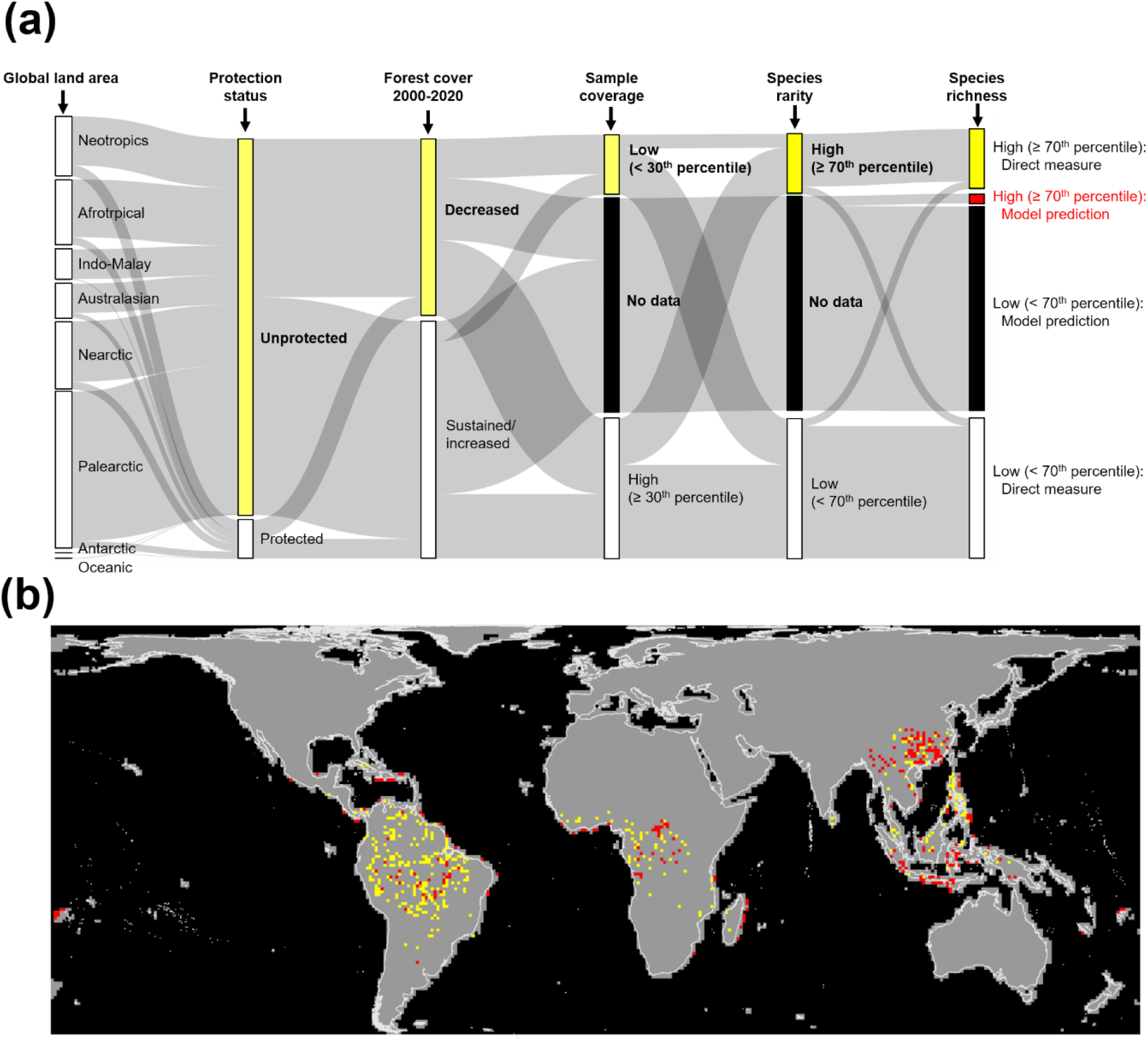
Spatial priority areas for improving sample completeness of species occurrence records of woody angiosperms. (a) Composition of the attributes used to define the priority areas. (b) The geographical map. Yellow color represents the priority areas defined as low sample coverage (< 30^th^ percentile of sample coverage values among the grid cells), high species rarity (≥ 70^th^ percentile of species rarity values among the grid cells), high species richness (≥ 70^th^ percentile of species richness values among the grid cells), unprotected and forest cover was lost during 2000-2020. Red color represents the grid cells with similar values except no occurrence data, but predicted as potentially high species richness (≥ 70^th^ percentile of species richness) by the Random Forest model.

To visualize the urgent needs for botanical exploration, we selected the grid-cells (100 km × 100 km) with lower sample coverage (< 30th percentile among the grid cells), higher species richness (≥ 70th percentile among the grid cells), and rarity (≥ 70th percentile among the grid cells) from the currently unprotected and deforested areas (Figs. 6 and S26). Those areas represent spatial priority areas of future inventories where immediate assessments and evidence-based conservation decisions would be needed. We found the priority areas in South America, Central, and part of West Africa, subtropical China, and Indomalayan islands (Fig. 6). These areas are spatially consistent with botanically unexplored areas (*59*) with poor mobilization of existing occurrence data (*60*) in the known global biodiversity hotspots (*61, 62*). Notably, the priority map reflected not only information gaps as reported by previous studies (5), but also needs for preferential sampling efforts (including mobilization of existing records) that are informative to effectively fill the knowledge gap in plant biodiversity (*62, 63*).

### Concluding remarks

This study evaluated the patterns of global woody angiosperm species diversities (Hill numbers at the order 0, 1 and 2) using rarefaction/extrapolation based on biodiversity estimation theory. The estimated diversity patterns and assessment of environmental drivers demonstrated that climatic factors mostly shape global-scale diversity gradients and anomalies through species sorting with latitude and longitude (*40, 64*). These diversity gradients and anomalies were also influenced by the combination of spatial extent and grain size in the diversity estimation, which is conceptually linked to local/regional species pool size. Importantly, this study confirmed the non-linear latitudinal and longitudinal diversity gradients that were determined by the relative effects of different climatic variables, including historical components, as recently argued in marine biodiversity patterns, e.g. bimodality with a tropical decline or truncated bimodality in response to paleo-/modern climatic changes (*65, 66*). The geographical distribution of climatically favorable areas (e.g., warm, non-arid and moderate seasonality), current and historical, for harboring woody angiosperms created a symmetric unimodal diversity pattern peaking at the equator in the Americas, bimodality peaking at both the equator and southern high latitudes in Europe-to-Africa, and asymmetric unimodality peaking at northern mid-latitudes in Asia-to-Australia. Moreover, longitudinal anomalous diversity patterns of woody angiosperms peaking at South America and East Asia support historical imprints of the origin of tropical and temperate biome and their expansion across continents (*67*).

Our results revealed that there are areas, both inside and outside the global protected area network, where botanical sampling efforts have been inadequate or non-existent. This incomplete data can result in spatial disparities between biodiversity and protected area networks (*68*). Our findings demonstrated that locations with a potentially high number of rare and diverse woody angiosperms are present in the data-deficient areas, suggesting a conservation gap of the current protected areas. Prioritizing the inventory of biodiversity in threatened and unprotected locations is a pressing issue in enhancing the effectiveness of protected area expansion (*69*) in the post-2020 biodiversity framework.

## Materials and Methods

### Woody angiosperm species data

We prepared a candidate species list of woody angiosperms (here defined as species having lignified stem tissues including trees, shrubs, lianas, bamboos, palms, and cacti) comprising all angiosperm species except exclusively herbaceous families such as Orchidaceae or Cyperaceae. The candidate species list contained 223,724 species. Based on information in the national floras, botanical literature and various databases (see table S1 in the Supplementary Material for the source list), we judged woody species by checking whether the original botanical literature include the following words: woody, tree, shrub, trunk, undershrub, semi-shrub, palm, or culm. We standardized the species names following the World Checklist of Vascular Plants (https://wcvp.science.kew.org/), and integrated subspecies and varieties into the parental binomials. The final list of confirmed woody angiosperms comprised 123,878 species from 6,844 genera and 296 families (table S2). Based on the species list, we retrieved 40,770,307 occurrence records from existing databases and literature (tables S1 and S3). We retained only records with precise location information (longitude/latitude coordinates or locality names with <100 km precision). We removed records that were suspected to be a result of artificial introduction by verification in national floristic lists. After the data cleaning processes, the data set comprised 13,959,780 occurrence records for 82,974 species (67% of known woody angiosperms).

We converted the species occurrence records into species incidence data at the scale of approximately 10 km × 10 km cells in the Behrmann projection. To calculate sample completeness and estimate diversity, we defined four coarser grids, with cells of 100 km × 100 km, 200 km × 200 km, 400 km × 400 km, and 800 km × 800 km resolution, respectively. Within each cell of those coarser grids, we counted the frequency of species incidence across 10 km × 10 km equal area grid cells. Due to the known limitations of the analytical framework of diversity estimation (*70*), we excluded the cells for which the sample size was deemed inadequate: observed number of species was less than 6, number of the sub-gridded cells where at least one incidence was recorded was less than 6, or total number of species incidence was equal to the number of singletons (i.e., there were no species recorded in more than one 10 km × 10 km grid cell).

### Environmental data

To assess environmental drivers of species diversity, we obtained climatic information for the present day and the Last Glacial Maximum (LGM) from WorldClim (*71*). As surrogates of historical climatic stability, we calculated absolute differences in annual mean temperature and annual precipitation between the present day and the LGM. Data for actual evapotranspiration, potential evapotranspiration and aridity index were obtained from the Global High-Resolution Soil-Water Balance dataset (*72, 73*). Elevation data at 15-arcseconds resolution were obtained from the Geospatial Information Authority of Japan (https://www.gsi.go.jp/kankyochiri/gm_global_e.html).

### Diversity estimation

We calculated incidence-based species diversities (diversity at *q* = 0, 1 and 2, corresponding to species richness, Shannon diversity and Simpson diversity, respectively) (*15*) in each coarse grid-cell. Given that the relationships between diversity and number of incidences were often non-saturated, we could not know the true diversity nor use asymptotic diversity values to fairly and reliably compare diversity among grid cells. Instead, we standardized the estimated diversity using a combination of rarefaction and extrapolation based on sample completeness (*15*). We used sample coverage in each grid cell as a measure of sample completeness; the sample coverage is the proportion of the number of detected incidences to total (detected + undetected) incidences in the grid cell (*14*). We used sample coverage values of doubled reference sample size to capture a wide range of species diversities as possible within a reliable extrapolation range (*74*), while we slightly mitigated this limitation for extrapolation by using the percentiles of sample coverage values among the grid cells to standardize diversity (*17*). This means, if a percentile is 40th, the standardized diversity of each grid cell was obtained by extrapolation to more than double its reference sample size for 40% of the grid cells; for the other 60% of the grid cells, the standardized diversity of each grid cell was obtained either by rarefaction or by extrapolation to less than double its reference sample size. By testing several levels of standardization (see supplementary text), we confirmed that the choice of level did not change the outcome of geographical pattern analyses (see below) unless levels of standardization that were too low (first percentile) or too large (e.g., 100th percentile or asymptotic diversity) were selected. The former is because, when data in all grids were rarefied to a low coverage value, only a few species would be involved and thus the true geographical pattern could not be detected. The reason for the latter is that asymptotic diversity (especially for *q* = 0) is typically subject to severe negative bias.

### Geographical pattern analyses

We mapped global woody plant diversity and drew latitudinal and longitudinal diversity gradients in three longitudinal (Americas, Europe-to-Africa, and Asia-to-Australia) and three latitudinal (northern extratropics, southern extratropics, and tropics) zones. For each latitudinal/longitudinal zone, we fitted a Loess (locally estimated scatterplot smoothing) regression curve (smoothing parameter alpha = 0.6) and then compared the shape of the curves.

To detect the predominant environmental drivers of species diversity at the global and regional scales, we conducted regression analyses using three modelling approaches: ordinary least squares (OLS), generalized additive model (GAM) and Random Forest. In all models, we set log-scaled species diversity as the response variable. As the explanatory variables, we used annual mean temperature (Bio 1), temperature seasonality (Bio 4), annual precipitation (Bio 12), precipitation seasonality (Bio 15), actual evapotranspiration (AET), potential evapotranspiration (PET), aridity index (AI), climatic difference between the LGM and present day for temperature (D_temp_) and precipitation (D_prec_), average elevation (Elv) and standard-deviation of elevation (Elv.sd). In a preliminary analysis, we confirmed that the full model (including all these explanatory variables) was the best model based on the Akaike Information Criterion.

Individual relationships between species diversity and the environmental variables were visualized by plotting predictive lines (curves) using partial residuals. The relative importance of the explanatory variables was evaluated for OLS and Random Forest, based on the coefficients of partial determination for OLS, and the mean squared error in out-of-bag data for Random Forest. The explanatory power of each model was evaluated using the coefficient of determination (*R*^2^). The predictive power was evaluated using root mean squared errors computed by 10-fold cross-validation. In the main text, we present the results for the Random Forest model, which showed the best overall performances among the three modelling approaches (see supplementary materials for the results of the OLS and GAM approaches).

#### Priority map for future sampling

We visualize priority areas for future inventory as strategic sampling efforts that enable us to effectively fill knowledge gaps of woody angiosperm diversity. Specifically, we superimposed sample coverage, standardized species richness (at sample coverage = 0.82), species rarity, deforestation trend, and protected areas at the scale of 100 km × 100 km grid cells. We defined the species rarity as the total number of unique (those that are detected only once in a grid cell) and duplicate (those that are detected twice in a grid cell) species (*14*). The information on protected areas was downloaded from the World Database on Protected Areas (https://www.protectedplanet.net/en), and then protected areas were judged when a grid-cell overlapped with the polygons belonging to the IUCN protected area categories I to VI. We computed the percentage loss/gain of forest coverage to the total land area in each grid cell between 2000 and 2020 using the GLAD Global Land Cover and Land Use Change dataset (*75*). We identified the priority areas as the grid-cells (100 km × 100 km) with lower sample coverage (< 0.52 or the 30th percentile among the grid cells) and higher species rarity (≥ 118 species or the 70th percentile among the grid cells): these priority areas were selected within currently unprotected and deforested grid-cells (Fig.S26), which would be more threatened by habitat loss or degradation (*58*).

### Software

All analyses were performed in the R statistical environment (ver. 4.1.1) (*76*) with the following packages: ‘rgbif’ (*77*) for downloading species occurrence records from GBIF; ‘iNEXT’ (*78*) for estimating and standardizing species diversity; ‘mgcv’ (*79*) for conducting GAM analysis; ‘ranger’ (*80*) for conducting Random Forest analysis; ‘pdp’ (*81*) for calculating partial dependence plots of Random Forest model; ‘spm’ (*82*) for cross-validation of Random Forest model.

## Supplementary Materials

**Supplementary Text**

**Figs. S1 to S26**

**Tables S1 to S4**

**References** (83 to 211)

