## supplementary materials for "Occurrence-based diversity estimation reveals macroecological and conservation knowledge gaps for global woody plants"

**This PDF file includes:**

Supplementary Text
Figs. S1 to S26
Tables S1 and S4
References (83 to 209)

**Other Supplementary Materials for this manuscript include the following:**

Tables S2 to S3

**Supplementary Text**

**1. Sample coverage levels of standardisation of species diversity**

To fairly compare diversity values among locations, we need to consider the spatial inequality of sample completeness (*6*). In the case of the global diversity of woody plants, the observed number of species showed strong correlation with the number of species incidences (Fig. S1), indicating that the spatial distribution of sample completeness contaminated the observed species diversity patterns. Because the relationships between observed species diversity and number of species incidences were often unsaturated in our dataset, we could not use the asymptotic diversity value to 'fairly' compare the diversities among grid cells (*15*). Instead, we applied a non-asymptotic approach whereby species diversity is standardised using rarefaction (interpolation) or extrapolation based on sample completeness (*14*).

Sample completeness can be evaluated using sample coverage, i.e. the proportion of the number of individuals (or frequency of incidences) detected in the focal assemblage. The sample coverage is more accurate than the conventional richness ratio (i.e. observed species richness/Chao-2 estimator), which is positively biased when sample size is inadequate (*14*). As a guideline, Chao and colleagues recommended using the minimum sample coverage value of doubled reference sample size for reliable extrapolation (*15*). However, in macro-scale studies, the magnitude of difference in true diversity could be huge ($10^1$ –$10^4$), and species-poor sites (e.g. temperate regions) tend to be explored better (higher sample coverage) than species-rich sites (e.g. tropical regions). Indeed, in our data set, sample coverage showed a large spatial variation, especially at the finest spatial resolution (Fig. S2). In such a case, if we applied an too small sample coverage, we would miss diversity gradients in regions showing less species richness. To deal with this, we relaxed the restriction to extrapolation by using percentiles of sample coverage values of doubled reference sample size as the level of standardization (*17*). When we used the *p*-th percentile as the level of standardization, extrapolation to more than double its reference sample size will be applied to *p*% of the grid cells, while rarefaction or extrapolation to less than double its reference sample size will be applied to (100 - *p*) % of the grid cells. There is no prescribed percentile value. Therefore, we set several percentiles (1st, 5th, 10th, 20th, 30th, 40th and 50th percentiles; see Table S4 for the specific values of sample coverage at each percentile) and assessed the impact of arbitrary choice of the level of standardization on descriptions, interpretations, and modelling of geographical diversity patterns. We also included the asymptotic diversity (i.e., assuming ∞ sampling effort or sample coverage = 1) for comparison.

*Impacts of sample coverage-based standardisation on geographical diversity patterns*

Standardisation based on sample coverage influenced the shape of latitudinal diversity gradients (LatDGs) differently for the three longitudinal zones: North-to-South Americas, Europe-to-Africa, and Asia-to-Australia (Fi.g S3). The shape of the LatDG was relatively robust to the level of sample coverage-based standardisation in the North-to-South Americas zone (Fig. S3). A relatively minor difference was observed in the steepness of the LatDG slope in the transition from tropical to temperate regions; the observed species richness tended to underestimate the slope owing to the undervalue in the tropical regions. In the Europe-to-Africa zone, although the LatDG was unclear for the observed species richness owing to large intra-regional variation in Africa and super-high representation in Europe (Fig. S3), sample coverage-based standardisation clarified consistently high species richness within the tropical zone and a decreasing trend from the equator to the northern pole. In the Asia-to-Australia zone, the observed LatDG peaked in the Southern Hemisphere (around the Tropic of Capricorn) and decreased towards the northern pole (Fig. S3); the sample coverage-based standardisation toned down the southern peak, and depicted alternative peak around the northern middle latitudes which represents the transition from tropical to temperate regions. Although the level of standardization affected the absolute values of diversity (i.e., the vertical position of LatDGs), they did not have a large influence on the shape of LatDGs.

Longitudinal diversity gradients (LonDGs) were also influenced by sample coverage-based standardisation, although they were less sensitive than LatDGs (Fig. S4). In the north extratropics, the standardised species richness became too low (almost zero) at the smallest level of standardization (1st percentile of sample coverage), obscuring the LonDG and generating a dip in the eastern margin (>100 °). In the tropics and the south extratropic, the standardised species richness was highest in the western parts (South America and/or South Africa) and decreased towards the east, whereas the observed species richness was over-represented in the eastern regions (Fig. S4).

*Impacts of sample coverage-based standardisation on environmental driver analysis*

We analysed relationships between the diversities and environmental variables using three regression models: ordinary least squares (OLS), generalised additive model (GAM), and random forest (RF). We used log-scaled species richness as the response variable, and 11 environmental variables as the explanatory variables associated with energy, water, their seasonality, topography, habitat heterogeneity, and historical climatic stability: mean annual temperature (Bio1), temperature seasonality (Bio4), annual precipitation (Bio12), precipitation seasonality (Bio15), actual evapotranspiration (AET), potential evapotranspiration (PET), aridity index (AI), average elevation (Elv), standard deviation of elevation (Elv.sd), and differences in temperature ($D_{temp}$) and precipitation ($D_{prec}$) between the Last Glacier Maxima and the present day.

The influence of sample coverage-based standardisation depends on the environmental variables and the modelling frameworks. In the OLS model, the linear relationship with the log-scaled species richness (residuals after controlling for the effect of the other explanatory factors) was maintained well for Bio 12 and AET (Fig. S5), whereas the reverse relationship was observed for Bio 4, Bio 15, PET, AI and $D_{temp}$. The effect size was magnified in the standardised species richness for Bio1 and Elv, but weakened for Elv.sd and $D_{prec}$.

In the GAM model, the response curves (with the effect of the other explanatory factors fixed as their respective means) to individual explanatory factors differed between the standardised and observed species richness, but were qualitatively similar among the sample coverage levels (Fig. S6). The curve for AET was the most robust to presence/absence of sample coverage-based standardisation; a consistent positive monotonic linear relationship was observed. Meanwhile, sample coverage-based standardisation was most influential on the relationships with Bio1, PET and Elv.sd. The observed (also asymptotic) species richness was high at extremely low Bio 1 and high PET (>2500 mm). This suggests that the observed diversity suffered from a multicollinearity associated with the biased geographical representation (e.g., higher latitudes). Such unrealistic patterns were not observed for the sample coverage-based standardised species richness.

In the RF model, the partial dependency curves were generally stable to presence/absence of sample coverage-based standardisation and its levels (Fig. S7). An exception was Bio 1 for which the partial dependency tended to be higher between 5 and 15 °C for the observed (and asymptotic) species richness, compared with that for the standardised diversities (Fig. S7). The species diversities were higher in regions with higher Bio 1, Bio 12, AET, AI, Elv and Elv.sd, lower Bio 4, Bio-15, $D_{temp}$ and $D_{prec}$, and intermediate PET.

We evaluated the potential impacts of sample coverage-based standardisation on relative importance of the explanatory variables using the coefficients of partial determination ($r^2$) and the mean squared error in out-of-bag data, for the OLS and RF models, respectively. The most important variable was consistently AET, regardless of sample coverage-based standardisation and the modelling approach (Figs. S8 and S9). In OLS, the model with the observed species richness tended to overvalue the relative importance of Bio 15, but undervalued Bio 1. In RF, the overall importance ranking was relatively stable against sample coverage-based standardisation, whereas the rank of $D_{temp}$ (historical temperature change) changed depending on the level of standardization: the importance increased with reduction in the level, suggesting that $D_{temp}$ would have better explanatory power when ignoring the diversity variation in species-poor regions, and may act as a dichotomous factor at the global scale (i.e., historically stable vs unstable sites).

The RF model showed the highest explanatory and prediction performance among the three modelling frameworks (Figs. S10 and S11). In all approaches, the explanatory power ($R^2$) was higher and the predictive error (root mean squared error based on a 10-fold cross-validation test) was smaller for the sample coverage-based standardised diversities than for the observed diversity. The explanatory and predictive performance was generally comparable among the level of standardisation, but slightly better in the intermediate levels (20th–40th percentiles).

Finally, we projected the regression models on the geographical space to visually check how the influence of sample coverage-based standardisation transmitted to spatial predictions. The spatial predictions based on the observed and asymptotic species richness were highly influenced by the geographical pattern of sample coverage (Figs. 1 and S12). The prediction based on low level standardisation (1st and 5th percentiles) failed to capture the diversity gradients in species-poor, higher-latitudinal regions. For the predictions based on the intermediate levels of sample coverage-based standardisations, the OLS model reflected a large-scale diversity trend from species-rich lower latitudes to species-poor higher latitudes, but obscured intraregional diversity variation. The nonlinear frameworks (GAM and RF) were successful in visualizing local diversity peaks (South America, central Africa, and south China) as well as the large-scale diversity trends from lower to higher latitudes (Fig. S12).

**2. Impacts of spatial resolution on species diversity analyses**

The sample coverage is dependent on spatial resolution (grid-cell size): the courser the spatial resolution, the higher the sample coverage on average, and the smaller its variance (Fig. S2). To test potential impacts of spatial resolution on the species richness estimations and biogeographical patterns, we repeated the above-mentioned suite of analyses at coarser spatial resolutions (~200 km × 200 km, 400 km × 400 km, and 800 km × 800 km).

Even at the courser spatial resolutions (≥ 200 km × 200 km), we observed LatDGs (Fig. S13). Species richness was highest in the tropics (the North-to-South Americas and the Europe-to-Africa zones) or the tropics–extratropics boundary (the Asia-to-Australia zone), and decreased toward higher latitudes. The LonDGs were relatively sensitive to the spatial resolution, especially for the north extratropics (Fig. S14). At the finest resolution (100 km × 100 km), species richness was higher in the eastern region (East Asia) than in the western regions (Europe and North America), whereas at the courser resolutions, a bimodal diversity pattern with comparable peaks in the eastern North America and East Asia were observed. In the tropics, at any spatial resolution, the species richness was highest in South America. In the southern extratropics, the species richness was highest between 45°W and 45°E (eastern South America and the tip of South Africa) at any spatial resolution. In LatDGs and LonDGs, the disparity between the observed (and asymptotic) and the sample coverage-based standardised species richness was greatest at the finest resolution (100 km × 100 km) but was reduced at the coarser resolutions.

Overall, the spatial resolution showed only marginal effects on the correlative relationships between species richness and the environmental variables (Figs. S15–S17), except for the environmental variables whose range was truncated at the coarser resolutions (Bio 12, Bio 15, AI, Elv.sd, and $D_{prec}$). In OLS models, the effect size and estimation error of regression coefficients tended to be larger at coarser resolutions in general (Fig. S15). In GAM models, the response curves tended to deviate among spatial resolutions, especially at the edges of variable ranges (Fig. S16). The shape of partial dependency plots in the RF models was relatively stable to the difference in the spatial resolution (Fig. S17).

AET was consistently the most important explanatory variable regardless of the spatial resolution, whereas the ranking of the other variables were depending on the spatial resolution (Figs. S8 and S9). The relative importance ranking (except for AET) was more sensitive to the spatial resolution in OLS (Fig. S8) than in RF models. In the RF models, the relative importance ranking, particularly for the four most important variables (AET, PET, Bio 1 and Bio 4) was stable (Fig. S9). Interestingly, at coarse resolutions (>400 km × 400 km), the relative importance of historical temperature change ($D_{temp}$) was higher in OLS and RF models (Figs. S8 and S9).

In general, the explanatory power ($R^2$) was slightly better, but the prediction error was greater, at coarser resolutions (Figs. S10 and S11). The RF models consistently showed the best explanatory and predictive performances across all spatial resolutions (Figs. S10 and S11). Exceptionally, the GAM models showed the highest $R^2$ at the 800 km × 800 km, which is likely to be an artefact of overfitting owing to a small sample size (*83*).

**3. Comparison of geographical patterns between diversities at different orders**

To test the potential influence of species incidence frequencies on biogeographical patterns, we checked the behaviour of species diversities at different orders. In a Hill number-based approach, species diversity ($D$) is represented by the following equation (24):

when $q \neq 1$, ${}^{q}D = \left(\sum_{i=1}^{S} p_i^q\right)^{1/(1-q)}$;

when $q = 1$, ${}^{1}D = exp\left(-\sum_{i=1}^{S} p_i \log p_i\right)$,

where $S$ is the total number of species in an assemblage, and $p_i$ is the relative abundance of the $i$-th species. The parameter $q$ controls the weighting for relative abundance: the larger $q$ is, the larger the weight for abundant (dominant) species (*15*). At $q = 0$, all species are treated equally, then ${}^{0}D$ represents number of species (or species richness); at $q = 1$, species are weighted by their relative abundance, then ${}^{1}D$ corresponds to the exponential of Shannon entropy; at $q = 2$, species are weighted by their squared relative abundance (i.e., relatively dominant species will receive a higher weight), then ${}^{2}D$ becomes the inverse of Simpson's diversity index. This formulation can be straightforwardly extended to species incidence data (*15*).

We compared the geographical patterns (LatDGs and LonDGs) and the results of environmental driver analysis between species richness ($q = 0$), Shannon diversity ($q = 1$) and Simpson diversity ($q = 2$) at the scale of 100 km × 100 km grid cells. To reduce the volume of supplementary materials, we only show the results of RF models for the observed and standardised (at the 40th percentile of sample coverage = 0.82) diversities.

The orders of diversity ($q = 0$, 1 and 2) did not change the biogeographical patterns of global woody angiosperm diversity, as shown in the global maps (Fig. S18). All diversities showed similar latitudinal and longitudinal gradients (Fig. S19 and S20). While the absolute values were different (${}^{0}D > {}^{1}D > {}^{2}D$), the patterns of partial dependency curves along the environmental gradients in RF models were similar between the orders (Fig. S21). The relative importance ranking of environmental variables in RF models was also similar among the orders (Fig. S22). Because the rarefaction/extrapolation estimators for high orders ($q > 0$) are nearly unbiased and valid for a wide range of prediction (*15*), the consistency of geographical patterns among different orders of diversities ($q = 0$: species richness, $q = 1$: Shannon diversity, $q = 2$: Simpson diversity) suggests a robustness of our findings.

**4. Regional difference in environmental drivers of species diversity**

To test the generality or region-specificity of environmental drivers of species richness, we conducted a regional-scale environmental driver analysis (with the RF approach) using the standardised species richness (at the 40th percentile of sample coverage = 0.82) at the scale of 100 km × 100 km grid cells. We used the three longitudinal (North-to-South Americas, Europe-to-Africa, and Asia-to-Australia) and three latitudinal (north extratropics, tropics, and south extratropics) zones as the unit of regions.

Consistent relationships among the regions were observed for AET (positive monotonic), and Bio 12, AI, and Elv.sd (positive saturation) (Fig. S23). For the other variables, the relationships differed among the latitudinal and longitudinal zones (Fig. S23). Along the temperature gradient (Bio 1), the standardised species richness was saturated in the south and north extratropics, South-to-North Americas, and Asia-to-Australia zones, whereas it steeply dipped at higher temperature in the tropics (>25 °C) and the Europe-to-Africa zone (>20 °C). Negative relationships with temperature seasonality (Bio 4) were evident in the North-to-South Americas and Europe-to-Africa zones, but not in the Asia-to-Australia, and longitudinal zones. Precipitation seasonality (Bio 15) showed high regional variation: unimodal patterns in the tropics, south extratropics, and Asia-to-Australia zone, a saturating pattern in the north extratropics, and a negative relationship in the Europe-to-Africa zone. PET showed a strong negative relationship in the tropics, a unimodal pattern in the North-to-South Americas zone, and a saturating pattern in the south extratropics; notably, the peak and saturating points were common among the regions (about 1500 mm). With regard to elevation (Elv), the standardised species richness increased rapidly from lowlands (Elv = 0 m) to several hundred metres in all regions except the north extratropics where a rapid increase was observed only above 1,000 m. Historical temperature change ($D_{temp}$) showed positive relationships within a small degree (<5 °C), but changed to negative with greater change in temperature; this trend was most prominent in the North-to-South Americas zone. Historical precipitation change ($D_{prec}$) showed an abrupt decline of diversity at ~1,000 mm in the tropics, and a weak negative relationship in the Asia-to-Australia zone, whereas no clear trends were observed in the other regions.

The relative importance ranking of the environmental variables differed among the zones (Fig. S24). For the LatDGs (North-to-South Americas, Europe-to-Africa, and Asia-to-Australia zones), energy (AET), the climatic seasonality (Bio 4 and Bio 15) and historical temperature change ($D_{temp}$) showed high relative importance. For the LonDGs (north extratropics, tropics, and south extratropics), the factors relevant to availability of energy and water (AET, PET, AI, Bio 1 and/or Bio 12) were the most important drivers of species richness.

**Fig. S1.**

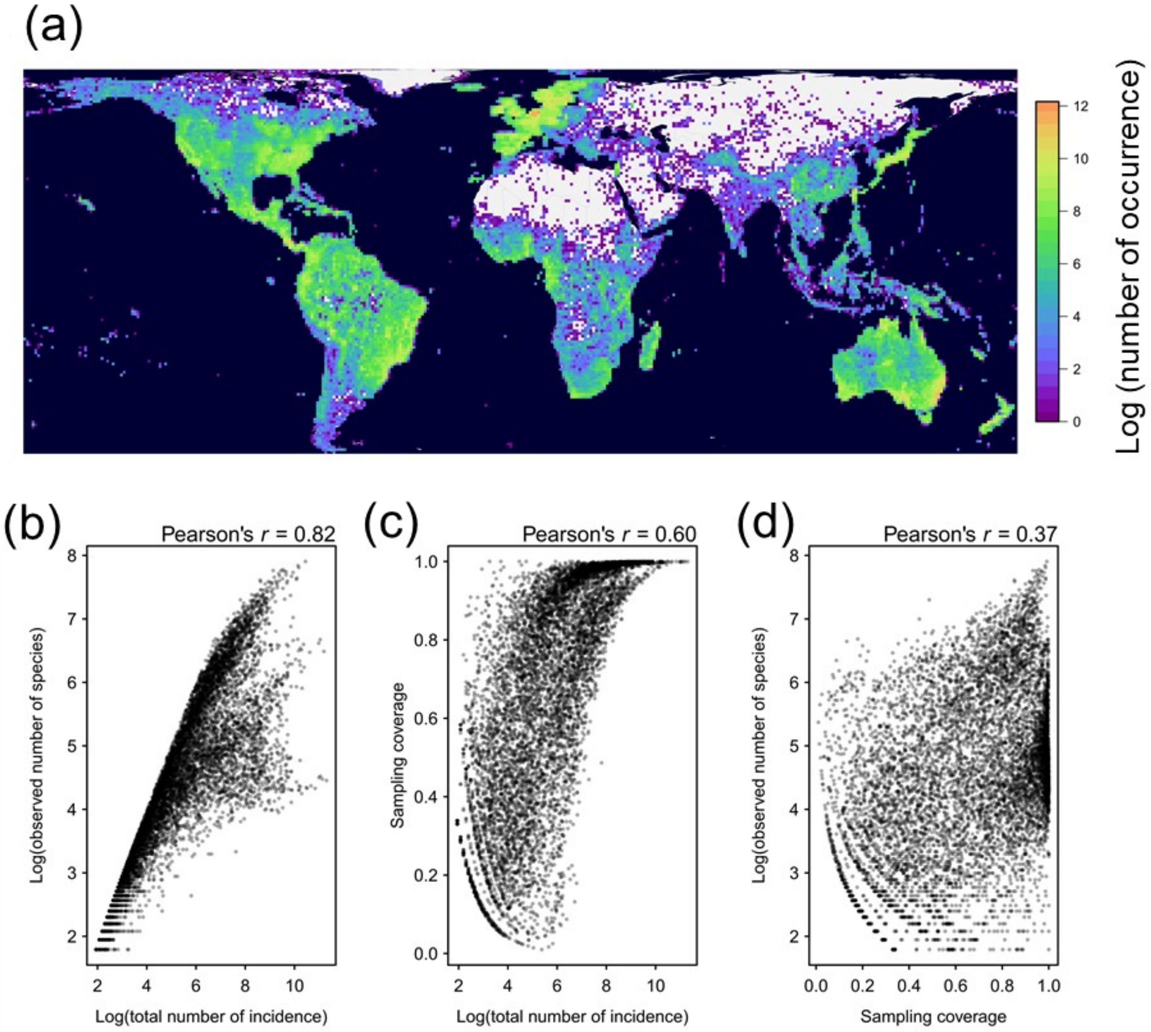


Global maps of log-scaled total number of occurrence records (a) and the relationships between observed number of species, total number of incidence, and sample coverage (b-d) at the level of 100 km × 100 km grid cells.

**Fig. S2.**

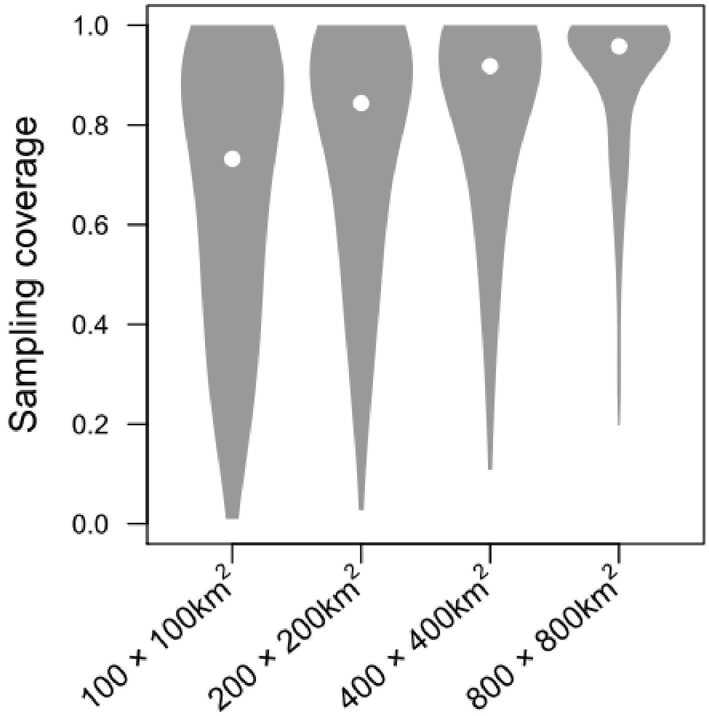


Violin plots of sample coverage values at four different spatial resolutions. White point shows mean value.

**Fig. S3.**

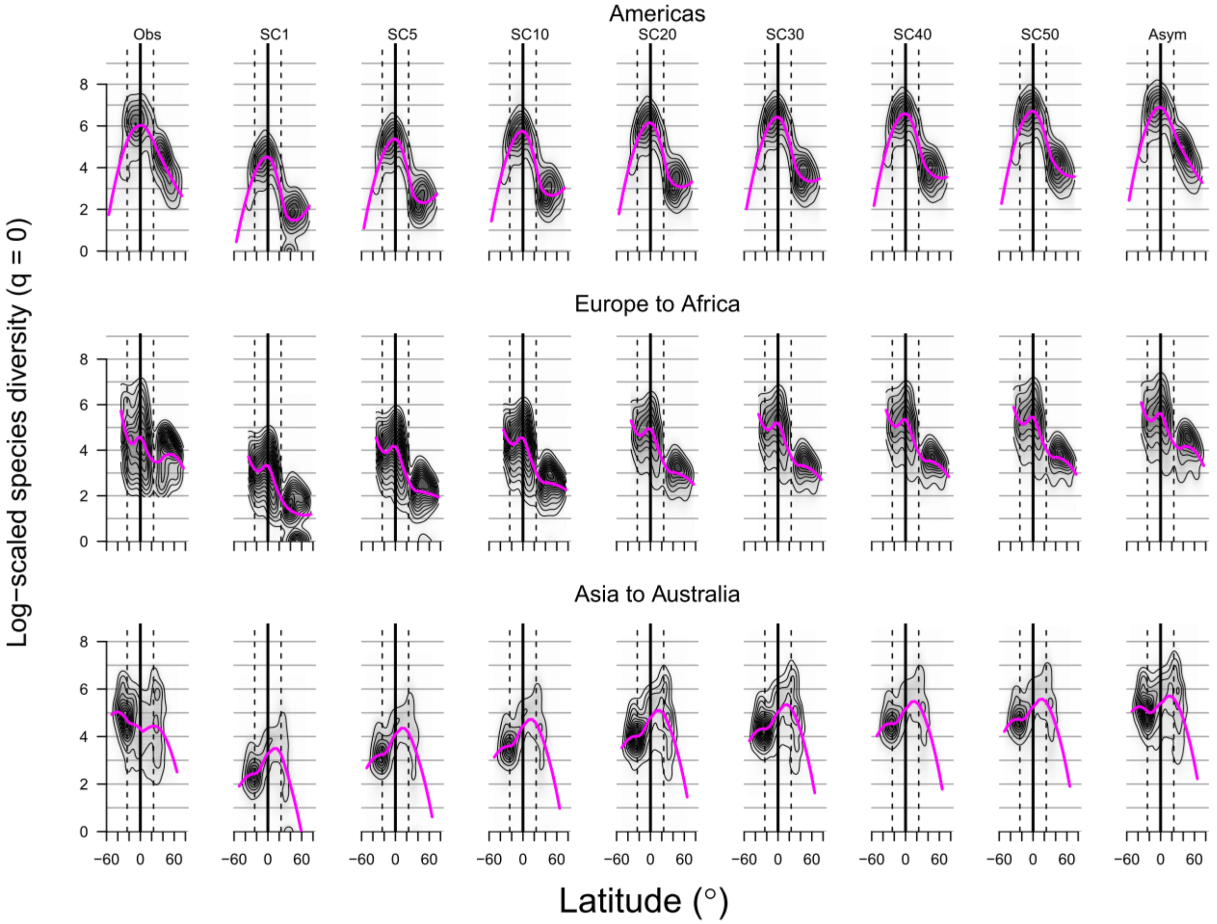


Latitudinal gradients of species richness at three longitudinal zones with different levels of standardization by sample coverage (SC): observed sample coverage (Obs), standardised at 1st, 5th, 10th, 20th, 30th, 40th and 50th percentiles of SC (SC1–50; see Table S4 for corresponding SC values), and asymptotic diversity (Asym; corresponding to SC = 100%). The diversity values are log-scaled. Loess (locally estimated scatterplot smoothing) curve (scaling parameter alpha = 0.6) is shown (pink line). Thick vertical line represents the equator. Dashed lines represent the tropics of Capricorn and Cancer.

**Fig. S4.**

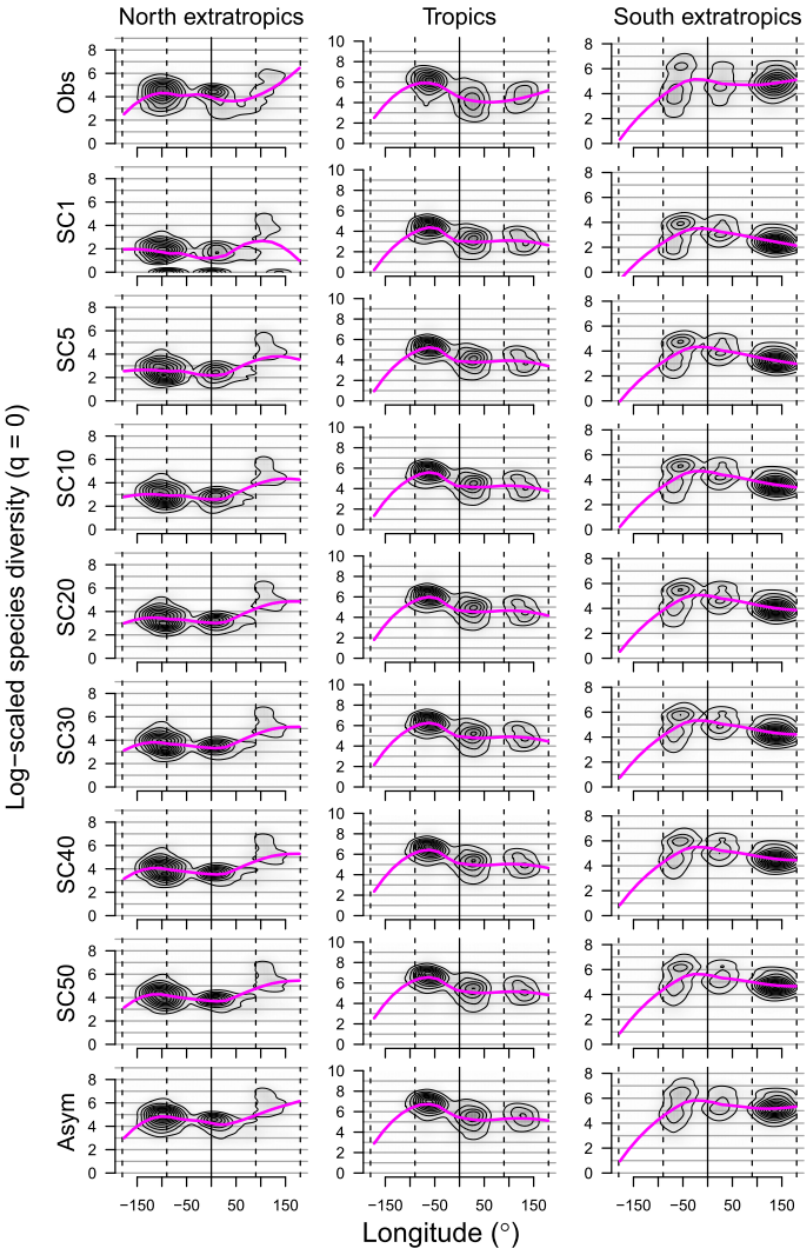


Longitudinal gradients of species richness at three latitudinal zones (tropics and north and south extratropics) with different levels of standardization by sample coverage (SC): observed sample coverage (Obs), standardised at 1st, 5th, 10th, 20th, 30th, 40th and 50th percentiles of SC (SC1–50; see Table S4 for corresponding SC values), and asymptotic diversity (Asym; corresponding to SC = 100%). The diversity values are log-scaled. Loess (locally estimated scatterplot smoothing) curve (scaling parameter alpha = 0.6) is shown (pink line).

**Fig. S5.**

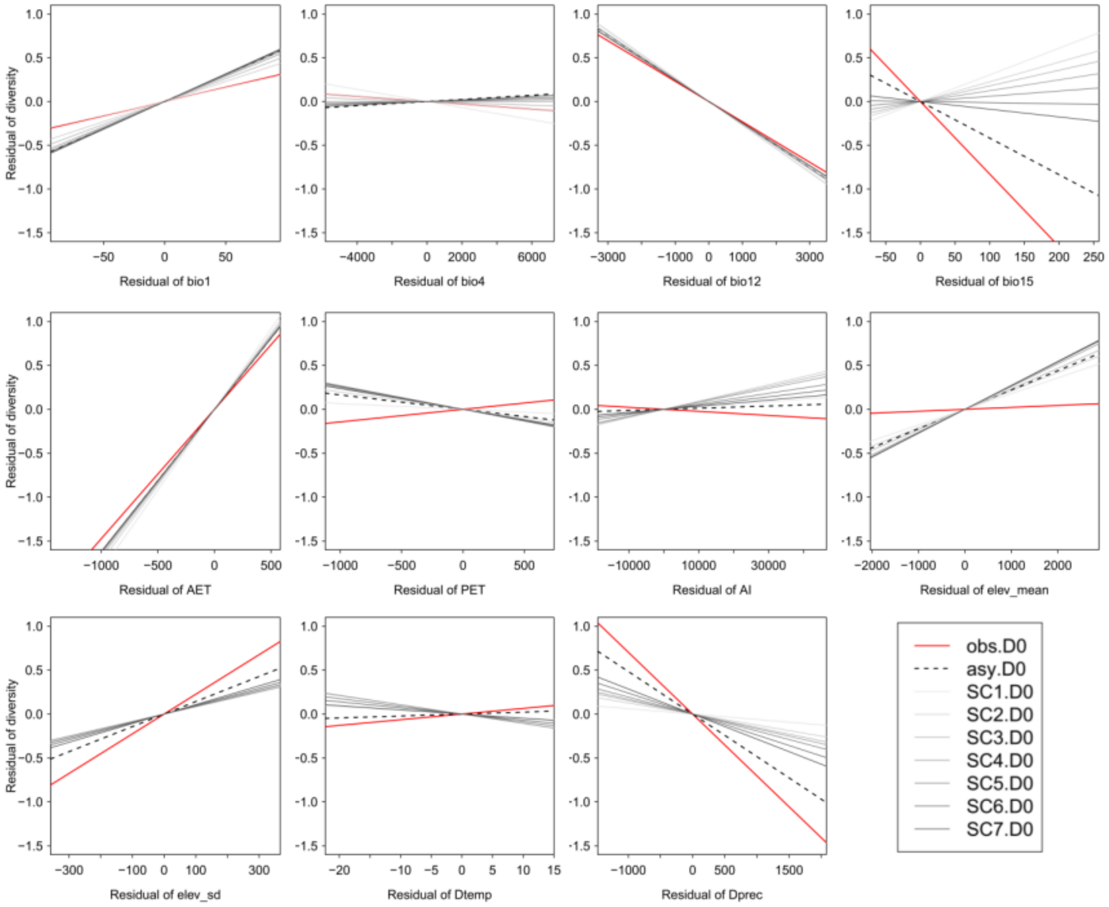


Linear relationship between species richness and environmental variables analysed by ordinary least squares model. The partial effect of each variable was evaluated by the residual regression. Species richness is standardised with different levels of sample coverage (SC): observed sample coverage (Obs), standardised at 1st, 5th, 10th, 20th, 30th, 40th and 50th percentiles of SC (SC1–50; see Table S4 for corresponding SC values), and asymptotic diversity (Asym; corresponding to SC = 100%). The environmental variables are mean annual temperature (Bio 1), temperature seasonality (Bio4), annual precipitation (Bio 12), precipitation seasonality (Bio 15), actual evapotranspiration (AET), potential evapotranspiration (PET), aridity index (AI), average elevation (Elv), standard deviation of elevation (Elv.sd), and differences in temperature ($D_{temp}$) and precipitation ($D_{prec}$) between the Last Glacier Maxima and the present.

**Fig. S6.**

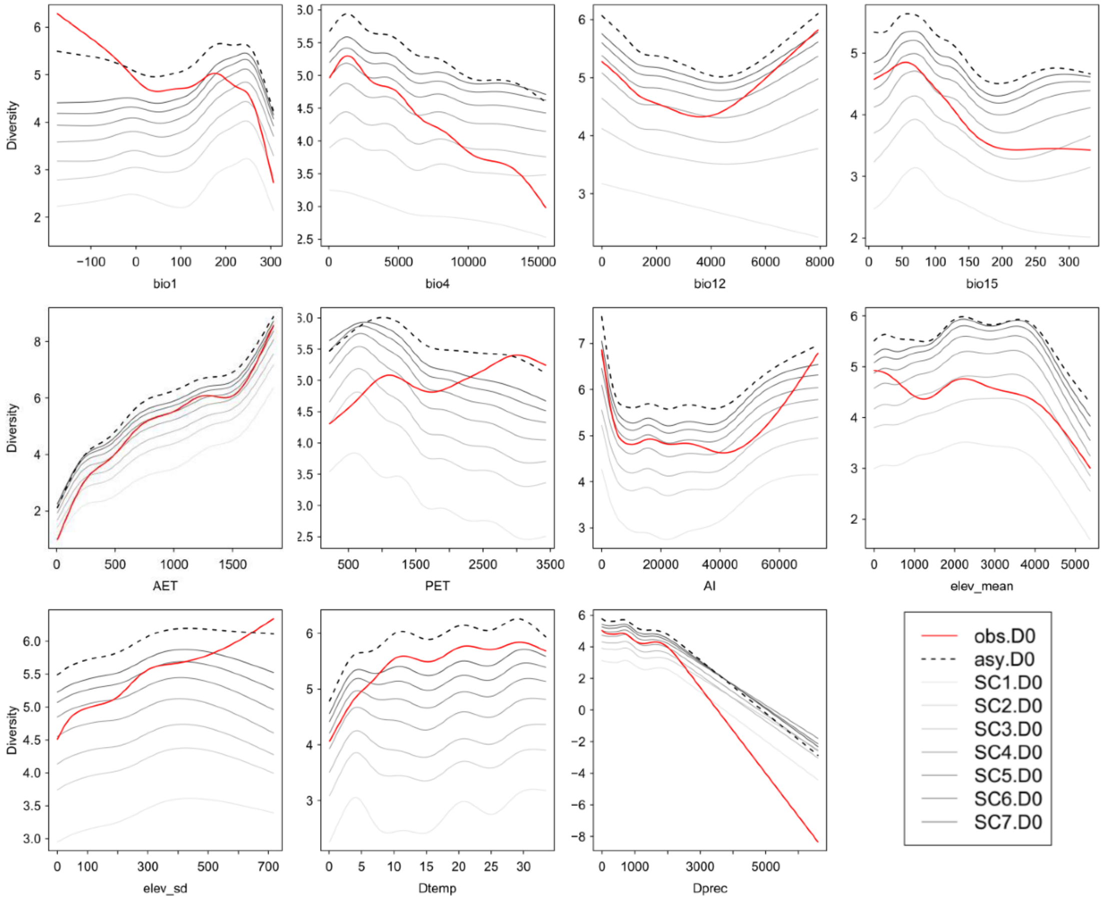


Relationship between species richness and environmental factors analysed by generalized additive model. Species richness is standardised with different levels of sample coverage (SC): observed sample coverage (Obs), standardised at 1st, 5th, 10th, 20th, 30th, 40th and 50th percentiles of SC (SC1–50; see Table S4 for corresponding SC values), and asymptotic diversity (Asym; corresponding to SC = 100%). The environmental variables are mean annual temperature (bio1), temperature seasonality (Bio 4), annual precipitation (Bio 12), precipitation seasonality (Bio 15), actual evapotranspiration (AET), potential evapotranspiration (PET), aridity index (AI), average elevation (Elv), standard deviation of elevation (Elv.sd), and differences in temperature ($D_{temp}$) and precipitation ($D_{prec}$) between the Last Glacier Maxima and the present.

**Fig. S7.**

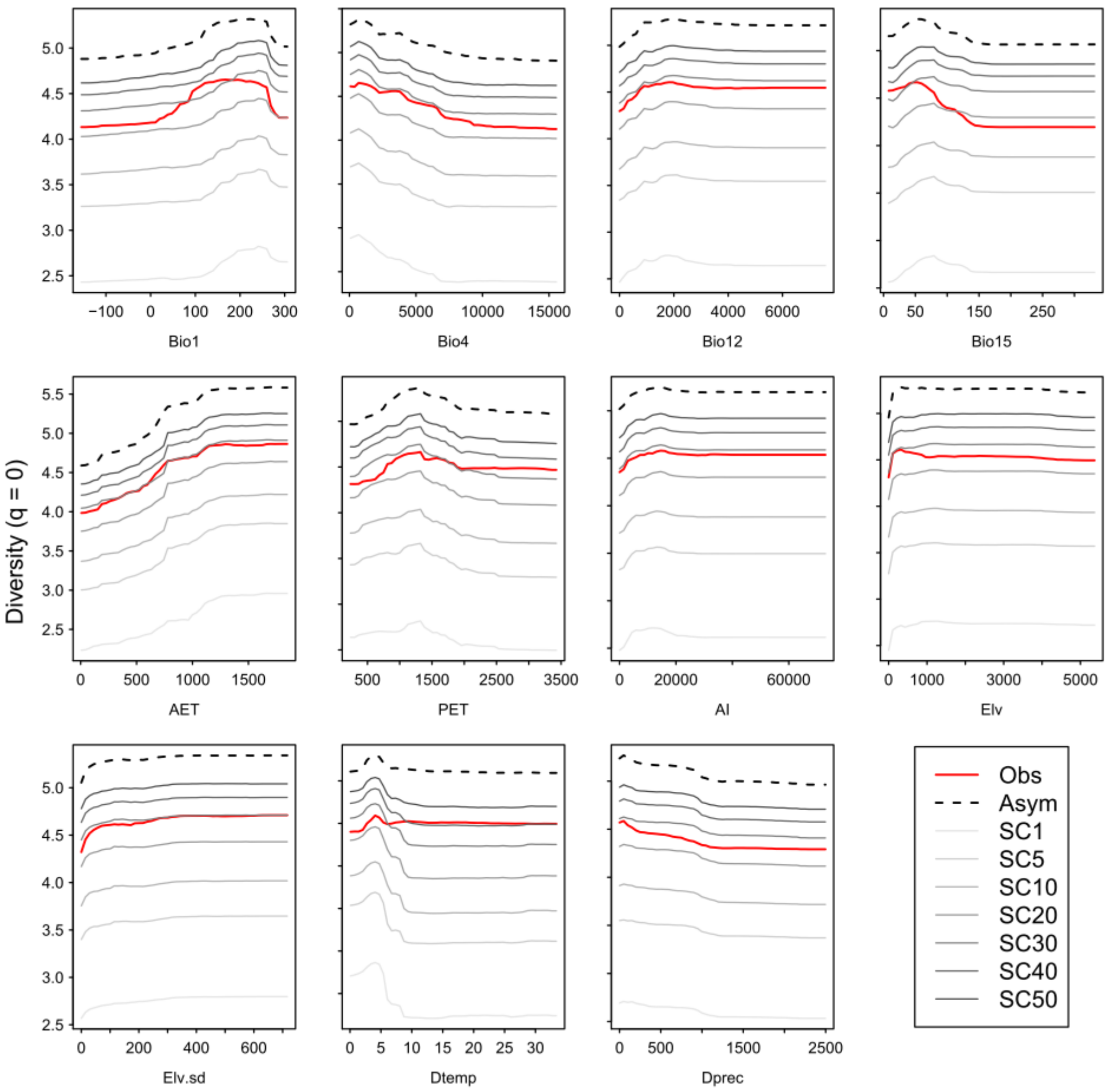


Partial dependency plot between species richness and environmental factors analysed by random forest model. Species richness is standardised with different levels of sample coverage (SC): observed sample coverage (Obs), standardised at 1st, 5th, 10th, 20th, 30th, 40th and 50th percentiles of SC (SC1–50; see Table S4 for corresponding SC values), and asymptotic diversity (Asym; corresponding to SC = 100%). The environmental variables are mean annual temperature (Bio 1), temperature seasonality (Bio 4), annual precipitation (Bio 12), precipitation seasonality (Bio 15), actual evapotranspiration (AET), potential evapotranspiration (PET), aridity index (AI), average elevation (Elv), standard deviation of elevation (Elv.sd), and differences in temperature ($D_{temp}$) and precipitation ($D_{prec}$) between the Last Glacier Maxima and the present.

**Fig. S8.**

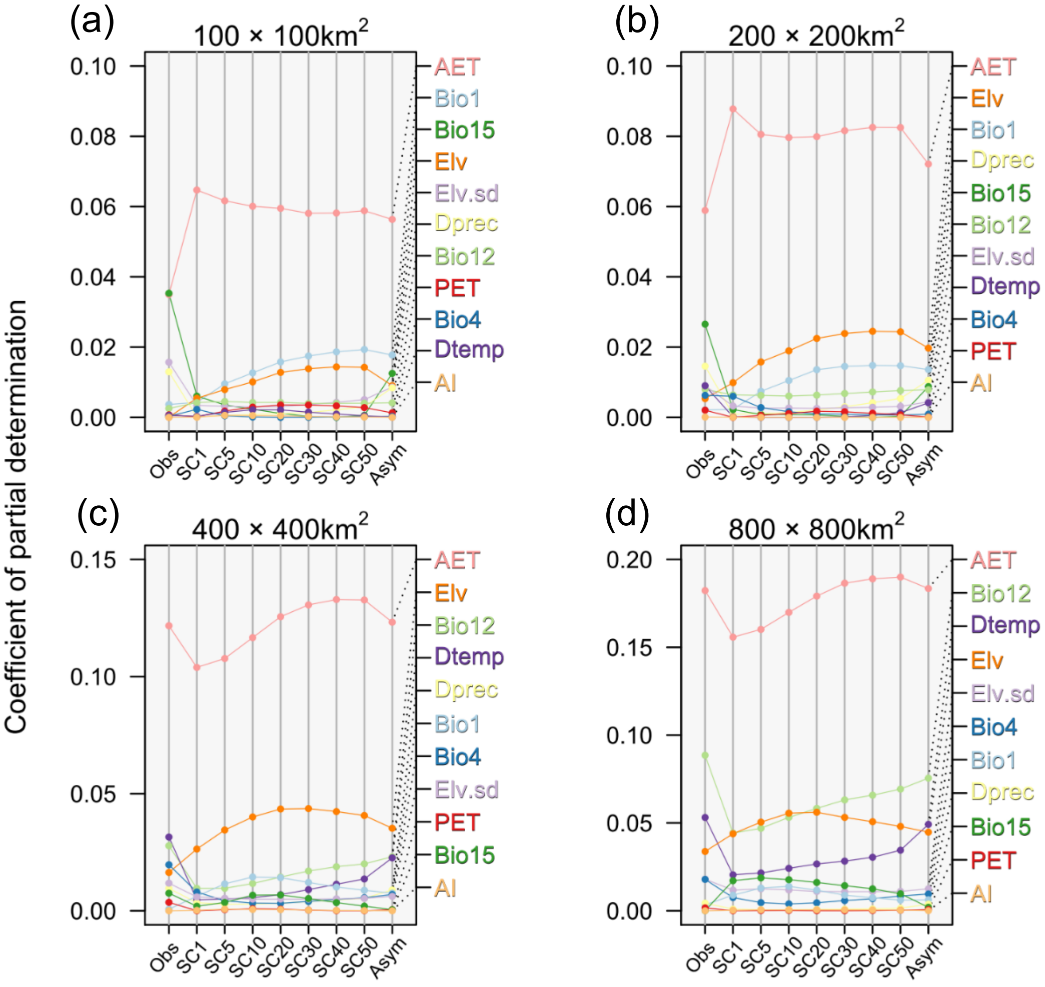


Relative importance (coefficients of partial determination) of environmental variables in the ordinary least squares model explaining species richness with different levels of standardization by sample coverage (SC): observed sample coverage (Obs), standardised at 1st, 5th, 10th, 20th, 30th, 40th and 50th percentiles of SC (SC1–50; see Table S4 for corresponding SC values), and asymptotic diversity (Asym; corresponding to SC = 100%) at different spatial resolutions. The environmental variables are mean annual temperature (Bio 1), temperature seasonality (Bio 4), annual precipitation (Bio 12), precipitation seasonality (Bio 15), actual evapotranspiration (AET), potential evapotranspiration (PET), aridity index (AI), average elevation (Elv), standard deviation of elevation (Elv.sd), and differences in temperature ($D_{temp}$) and precipitation ($D_{prec}$) between the Last Glacier Maxima and the present.

**Fig. S9.**

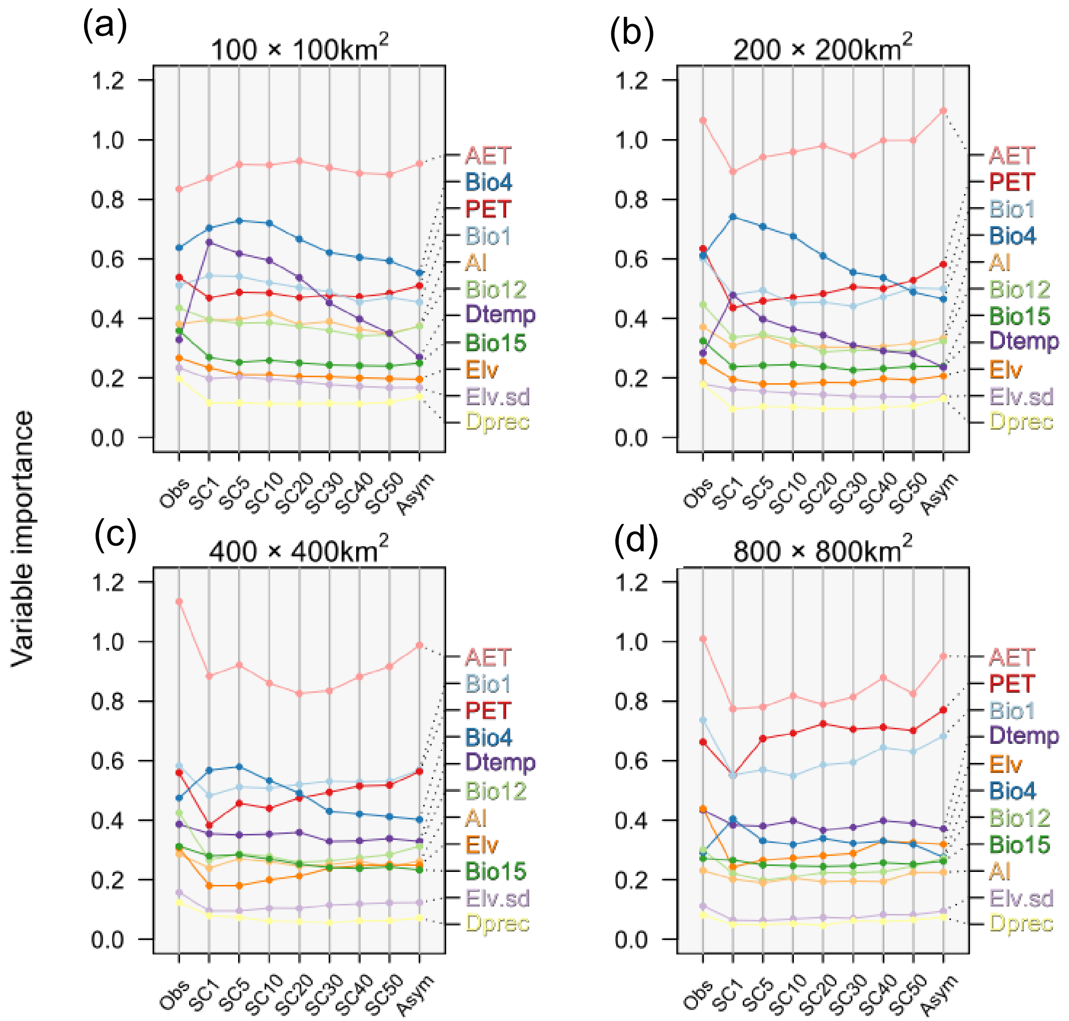


Relative importance of environmental variables in the random forest models explaining species richness with different levels of standardization by sample coverage (SC): observed sample coverage (Obs), standardised at 1st, 5th, 10th, 20th, 30th, 40th and 50th percentiles of SC (SC1–50; see Table S4 for corresponding SC values), and asymptotic diversity (Asym; corresponding to SC = 100%) at different spatial resolutions. The environmental variables are mean annual temperature (Bio 1), temperature seasonality (Bio 4), annual precipitation (Bio 12), precipitation seasonality (Bio 15), actual evapotranspiration (AET), potential evapotranspiration (PET), aridity index (AI), average elevation (Elv), standard deviation of elevation (Elv.sd), and differences in temperature ($D_{temp}$) and precipitation ($D_{prec}$) between the Last Glacier Maxima and the present.

**Fig. S10.**

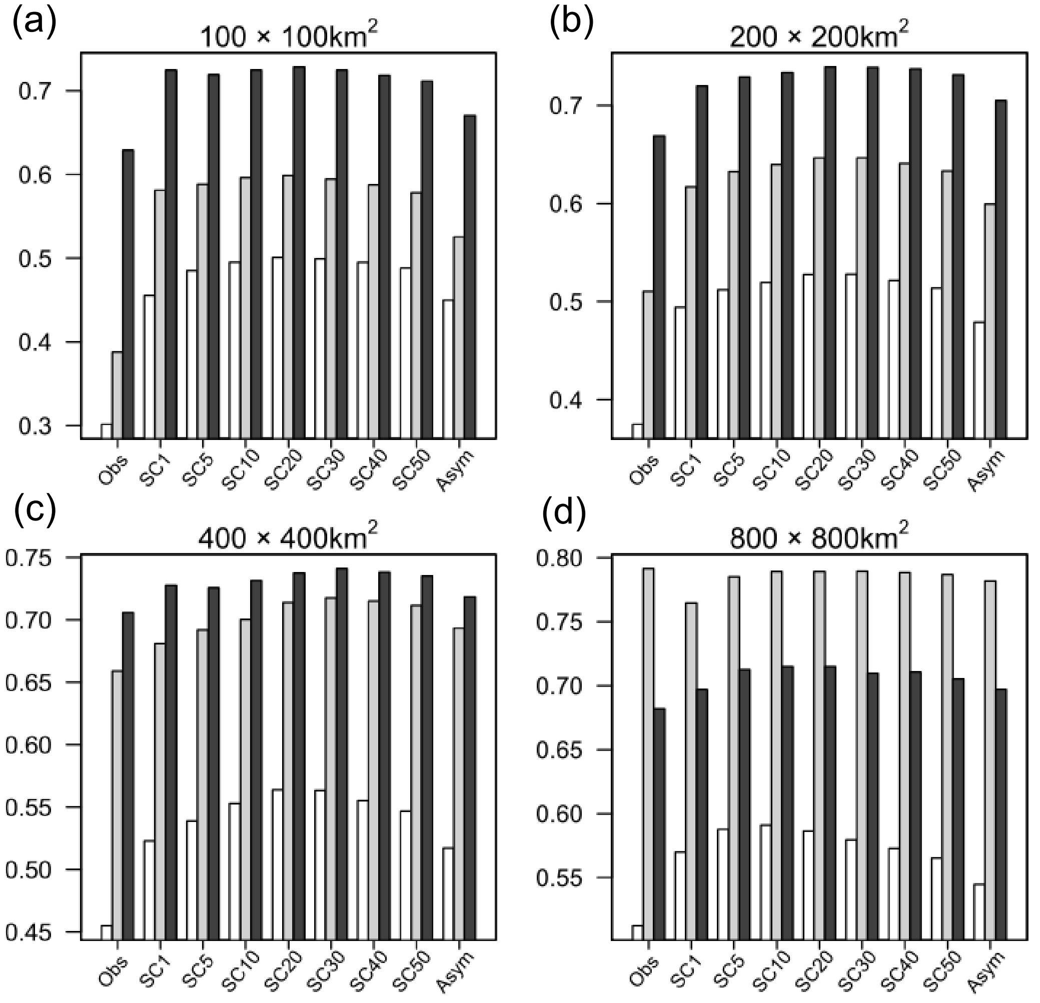


Explanatory power ($R^2$) of regression models (ordinary least squares = white, generalized additive model = light grey, random forest = dark grey) at different spatial resolutions explaining the relationship between environmental variables and species richness with different levels of standardization by sample coverage (SC): observed sample coverage (Obs), standardised at 1st, 5th, 10th, 20th, 30th, 40th and 50th percentiles of SC (SC1–50; see Table S4 for corresponding SC values), and asymptotic diversity (Asym; corresponding to SC = 100%).

**Fig. S11.**

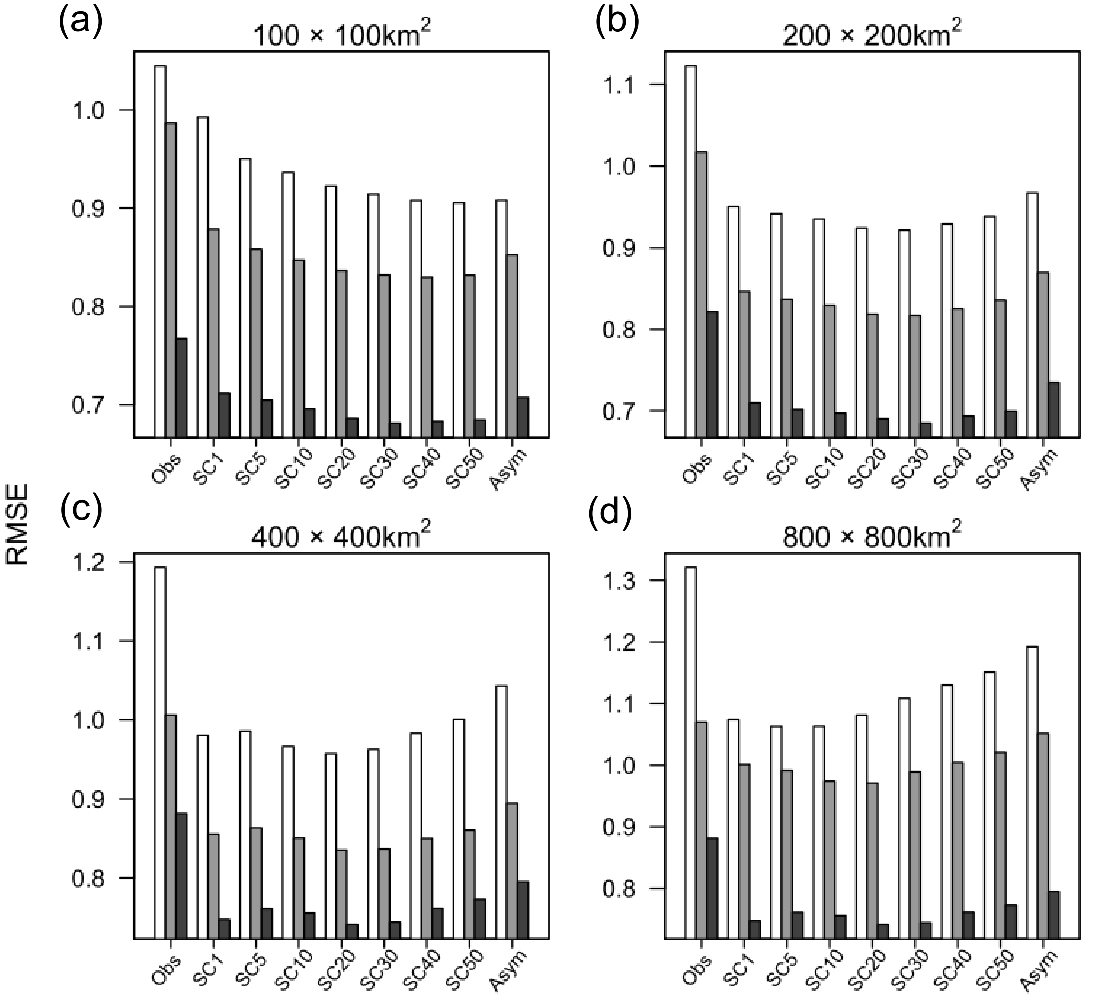


Predictive performance (root mean squared error from 10-fold cross-validation) of regression models (ordinary least squares = white, generalized additive model = light grey, random forest = dark grey) at different spatial resolutions, explaining the relationship between environmental variables and species richness with different levels of standardization by sample coverage (SC): observed sample coverage (Obs), standardised at 1st, 5th, 10th, 20th, 30th, 40th and 50th percentiles of SC (SC1–50; see Table S4 for corresponding SC values), and asymptotic diversity (Asym; corresponding to SC = 100%).

**Fig. S12.**

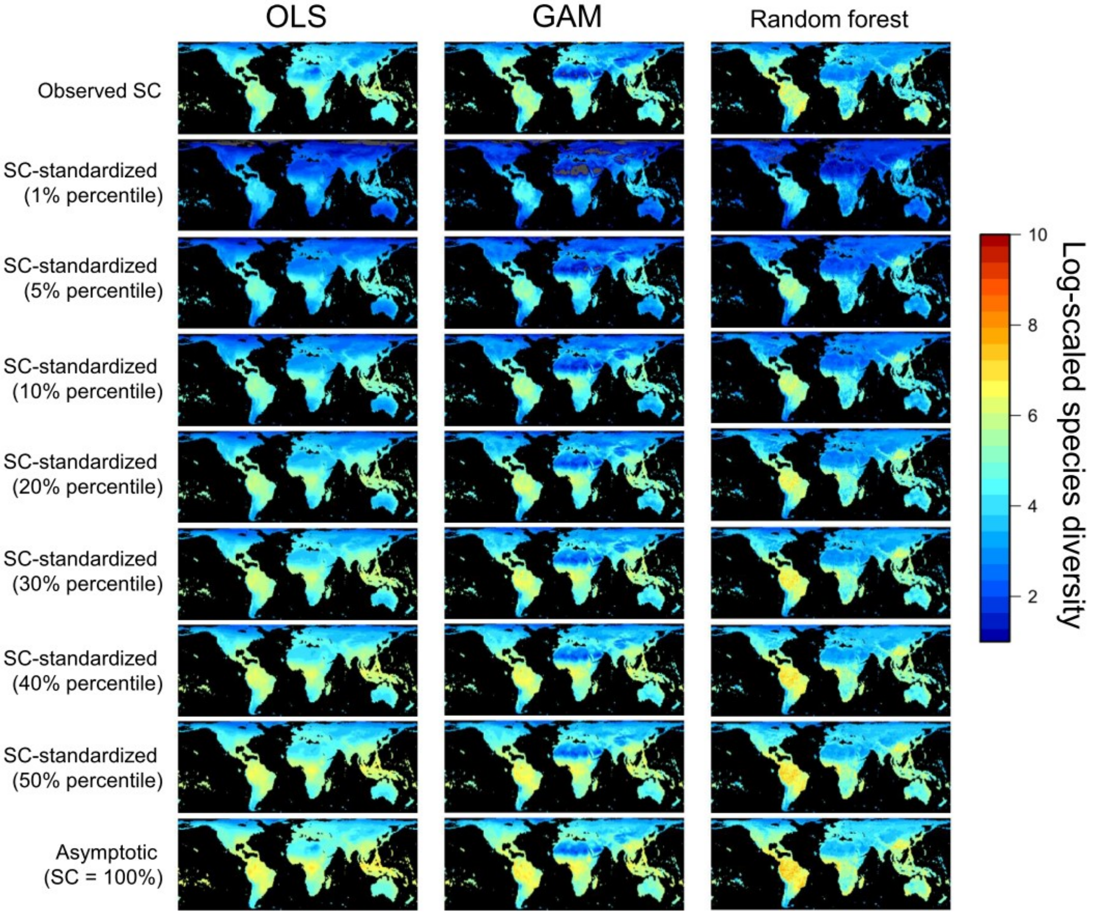


Projection of species richness with different levels of standardization by sample coverage (SC; see Table S4 for corresponding SC values) predicted using ordinary least squares (OLS), generalized additive model (GAM) and random forest with environmental variables at 100 km × 100 km grid cell level.

**Fig. S13.**

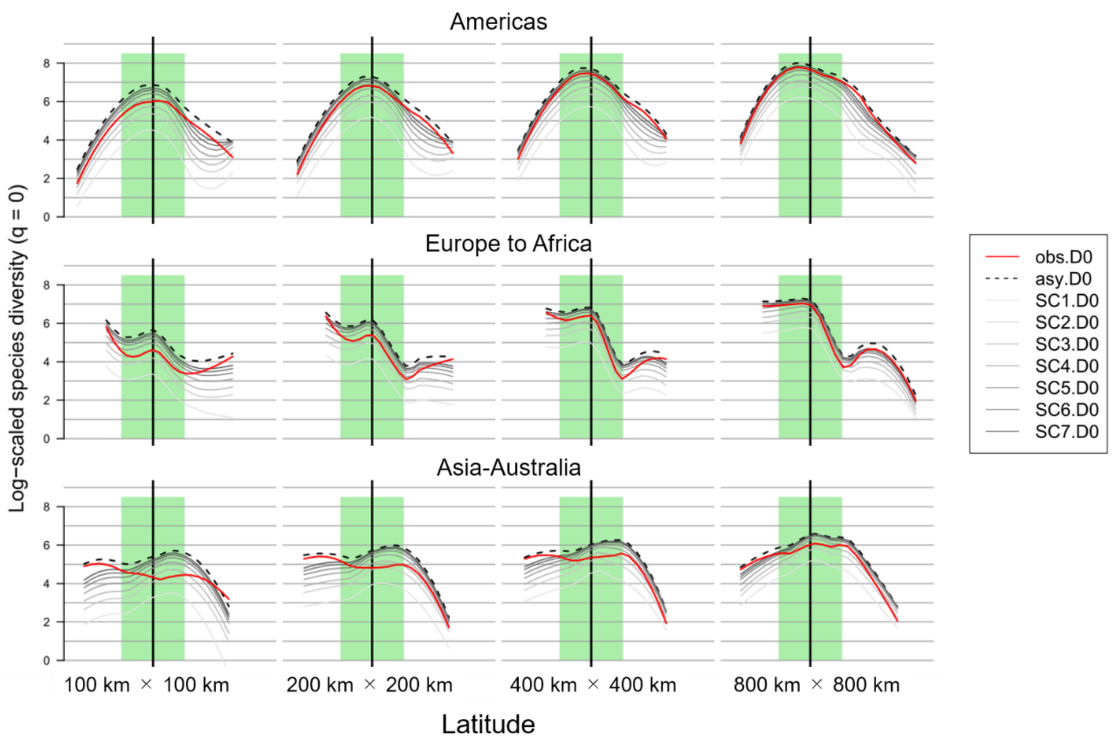


Latitudinal gradient of species richness at different spatial resolutions for three longitudinal bands: (a) the North-to-South Americas, (b) the Europe-to-Africa, and (c) the Asia-to-Australia zones. Loess (locally estimated scatterplot smoothing) curves (scaling parameter alpha = 0.6) are shown per species richness estimate with different levels of standardization by sample coverage (SC): observed sample coverage (Obs), standardised at 1st, 5th, 10th, 20th, 30th, 40th and 50th percentiles of SC (SC1–50; see Table S4 for corresponding SC values), and asymptotic diversity (Asym; corresponding to SC = 100%). The diversity values are log-scaled. Thick vertical line represents the equator. Shaded area represents the tropical zone between the tropics of Capricorn and Cancer.

**Fig. S14.**

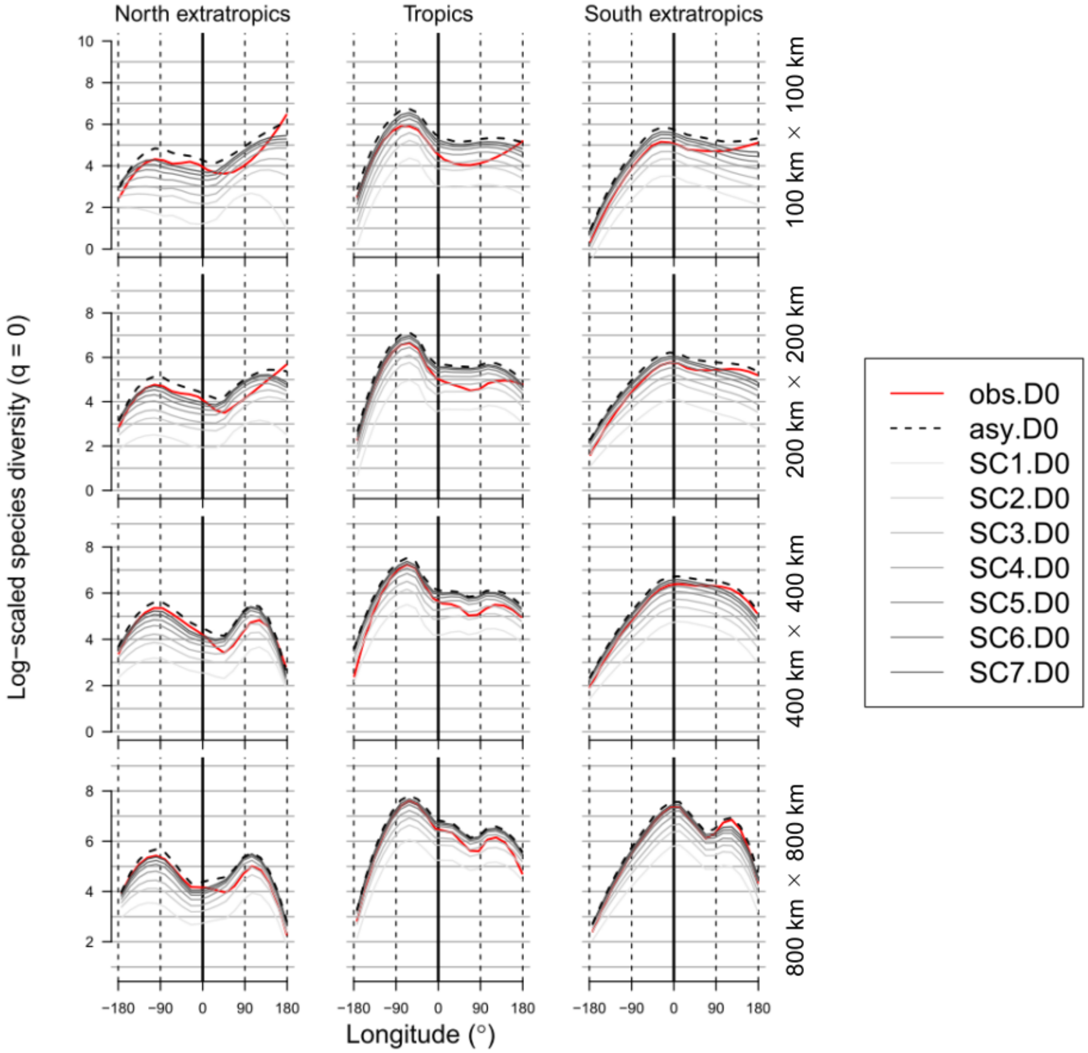


Longitudinal gradients of species richness at different spatial resolutions for three latitudinal zones (tropics and north and south extratropics) with different levels of standardization by sample coverage (SC): observed sample coverage (Obs), standardised at 1st, 5th, 10th, 20th, 30th, 40th and 50th percentiles of SC (SC1–50; see Table S4 for corresponding SC values), and asymptotic diversity (Asym; corresponding to SC = 100%). The diversity values are log-scaled. Loess (locally estimated scatterplot smoothing) curves (scaling parameter alpha = 0.6) are shown per species richness estimate with different levels of standardization by sample coverage (SC): observed sample coverage (Obs), standardised at 1st, 5th, 10th, 20th, 30th, 40th and 50th percentiles of SC (SC1–50; see Table S4 for corresponding SC values), and asymptotic diversity (Asym; corresponding to SC = 100%).. Thick vertical line represents the Prime meridian.

**Fig. S15.**

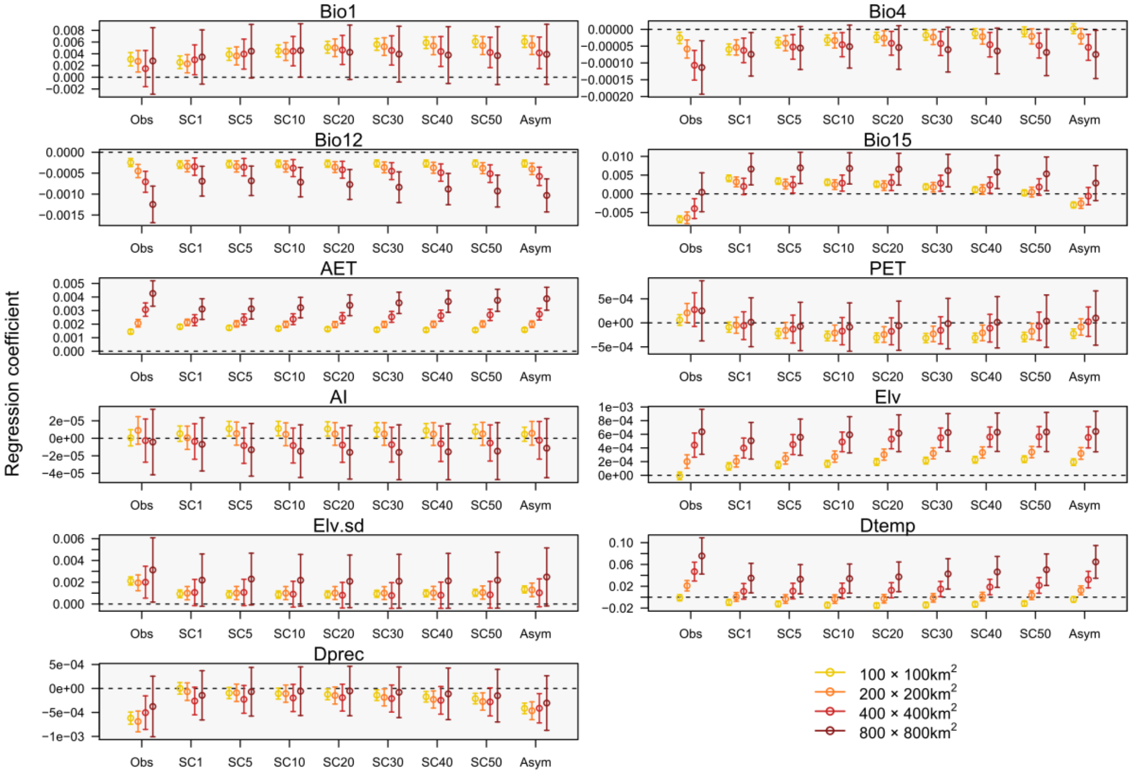


Impact of spatial-resolutions on the standardised regression coefficient of the environmental variables in ordinary least square regression model explaining species richness with different levels of standardization by sample coverage (SC): observed sample coverage (Obs), standardised at 1st, 5th, 10th, 20th, 30th, 40th and 50th percentiles of SC (SC1–50; see Table S4 for corresponding SC values), and asymptotic diversity (Asym; corresponding to SC = 100%) analysed using generalized additive model. The environmental variables are mean annual temperature (Bio 1), temperature seasonality (Bio 4), annual precipitation (Bio 12), precipitation seasonality (Bio 15), actual evapotranspiration (AET), potential evapotranspiration (PET), aridity index (AI), average elevation (Elv), standard deviation of elevation (Elv.sd), and differences in temperature ($D_{temp}$) and precipitation ($D_{prec}$) between the Last Glacier Maxima and the present.

**Fig. S16.**

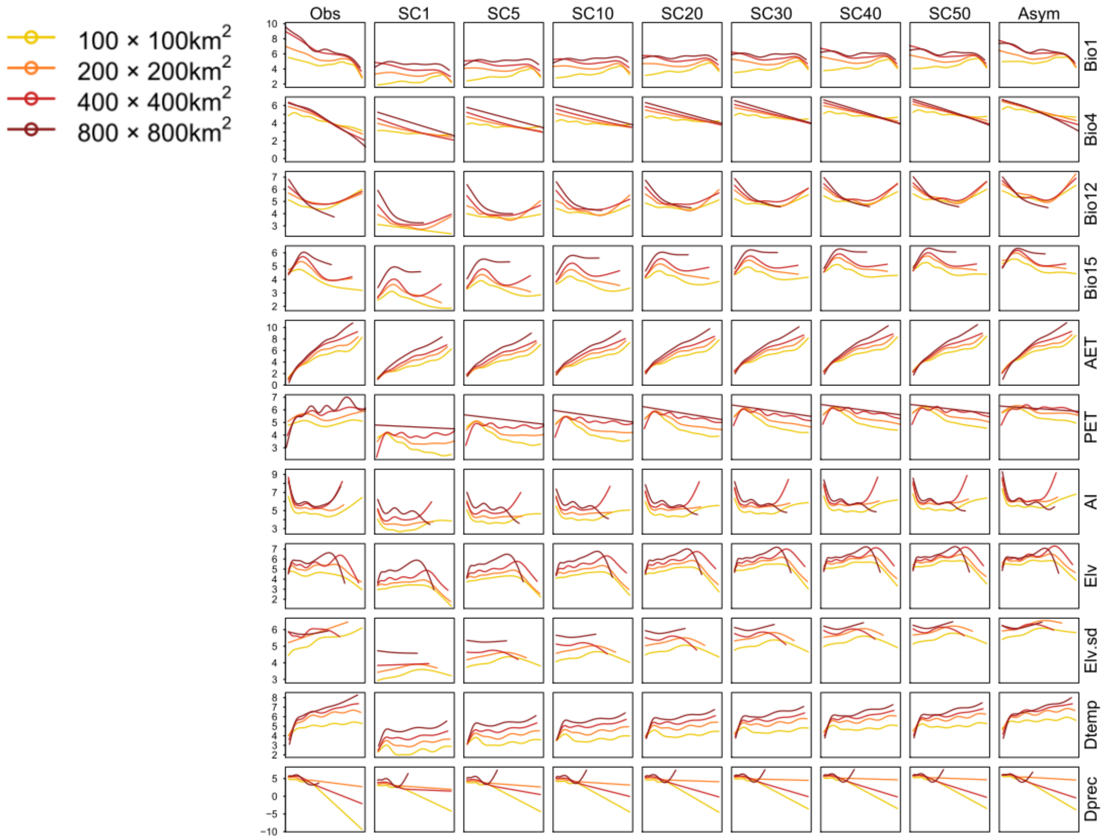


Impact of spatial-resolutions on the relationships between the environmental variables and species richness with different levels of standardization by sample coverage (SC): observed sample coverage (Obs), standardised at 1st, 5th, 10th, 20th, 30th, 40th and 50th percentiles of SC (SC1–50; see Table S4 for corresponding SC values), and asymptotic diversity (Asym; corresponding to SC = 100%) analysed using generalized additive model. The environmental variables are mean annual temperature (Bio 1), temperature seasonality (Bio 4), annual precipitation (Bio 12), precipitation seasonality (Bio 15), actual evapotranspiration (AET), potential evapotranspiration (PET), aridity index (AI), average elevation (Elv), standard deviation of elevation (Elv.sd), and differences in temperature ($D_{temp}$) and precipitation ($D_{prec}$) between the Last Glacier Maxima and the present.

**Fig. S17.**

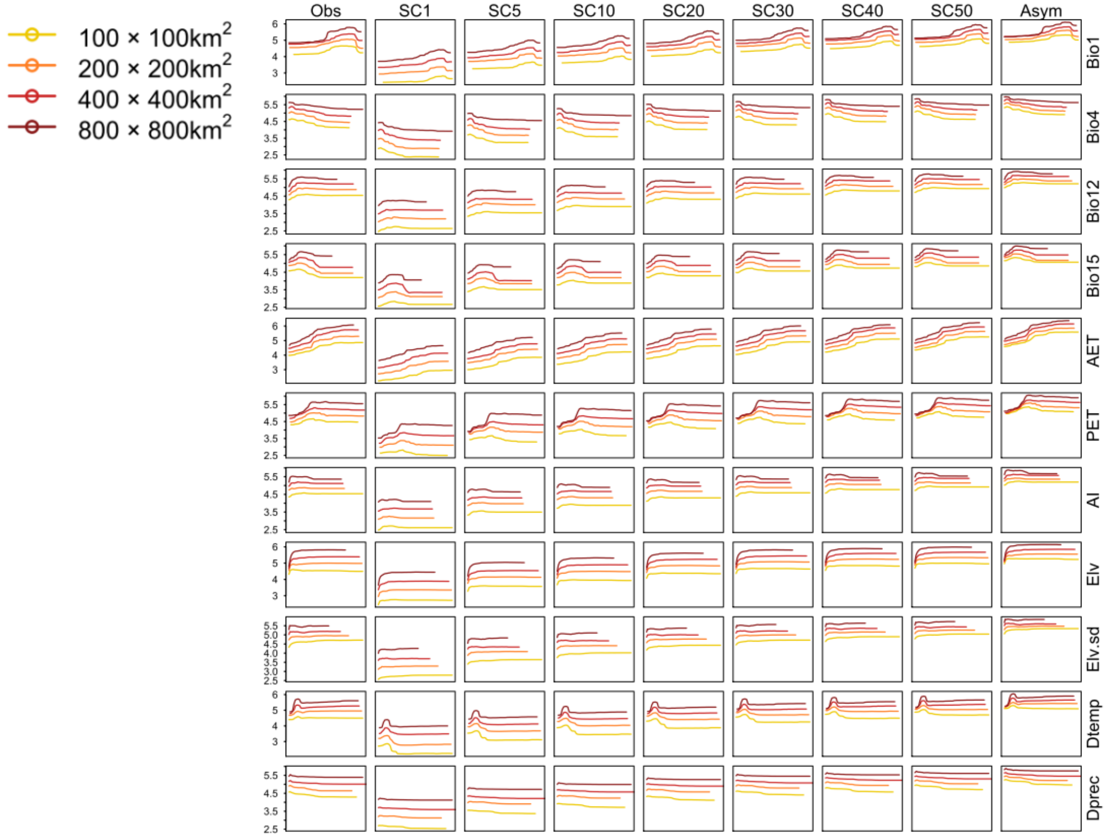


Impact of spatial-resolutions on the relationships between the environmental variables and species richness with different levels of standardization by sample coverage (SC): observed sample coverage (Obs), standardised at 1st, 5th, 10th, 20th, 30th, 40th and 50th percentiles of SC (SC1-50; see Table S4 for corresponding SC values), and asymptotic diversity (Asym; corresponding to SC = 100%) analysed using random forest model. The environmental variables are mean annual temperature (Bio 1), temperature seasonality (Bio 4), annual precipitation (Bio 12), precipitation seasonality (Bio 15), actual evapotranspiration (AET), potential evapotranspiration (PET), aridity index (AI), average elevation (Elv), standard deviation of elevation (Elv.sd), and differences in temperature ($D_{temp}$) and precipitation ($D_{prec}$) between the Last Glacier Maxima and the present.

**Fig. S18.**

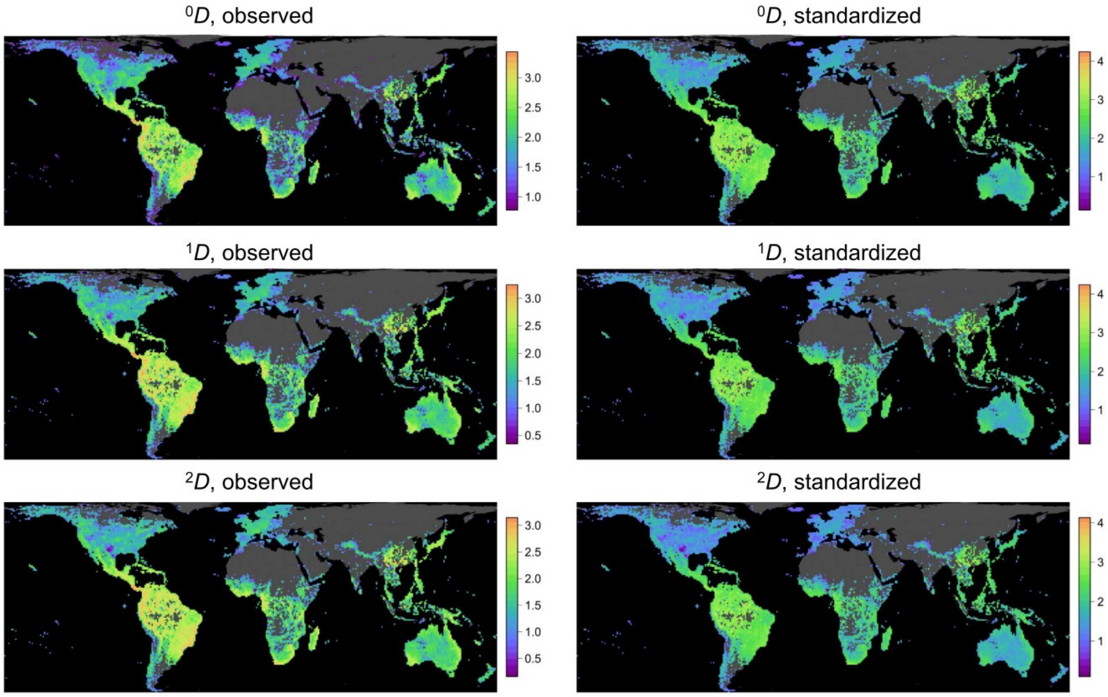


Geographical maps for the observed and standardised (at 40th percentile of sample coverage = 0.82) species diversity based on Hill numbers: species richness ($^{0}D$), Shannon diversity ($^{1}D$), and Simpson diversity ($^{2}D$), at approximately 100 km × 100 km grid cell level.

**Fig. S19.**

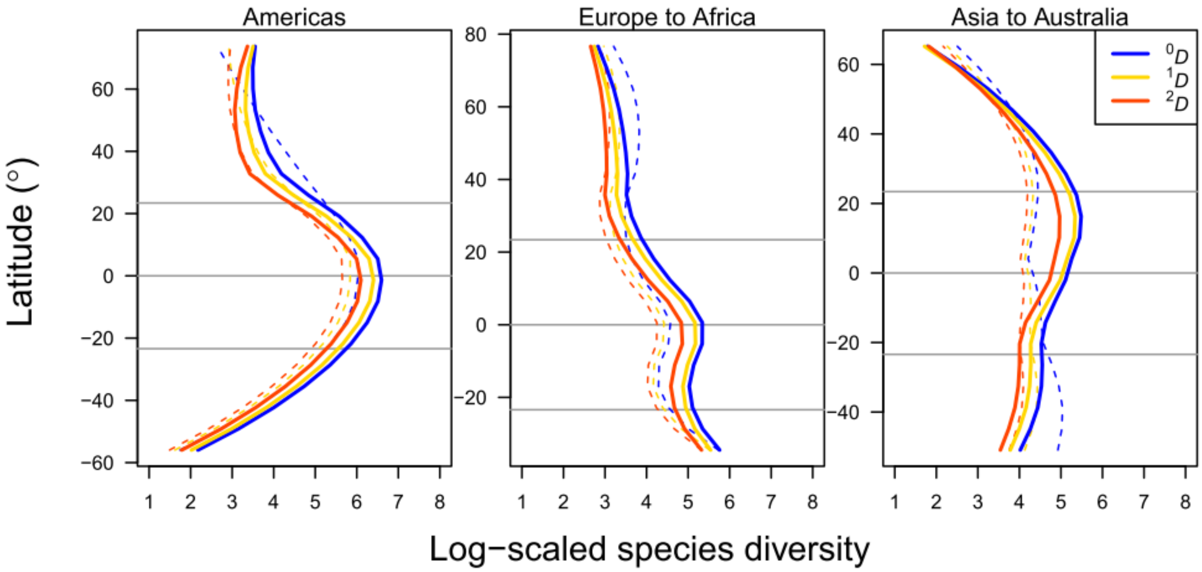


Latitudinal gradient of species diversity based on Hill numbers ($^qD$; $q$ = 0, 1, 2) at approximately 100 km × 100 km grid cell level. Dashed and solid lines represent observed and standardised (at 40th percentile of sample coverage = 0.82) values, respectively. Loess (locally estimated scatterplot smoothing) curves (scaling parameter alpha = 0.6) are shown.

**Fig. S20.**

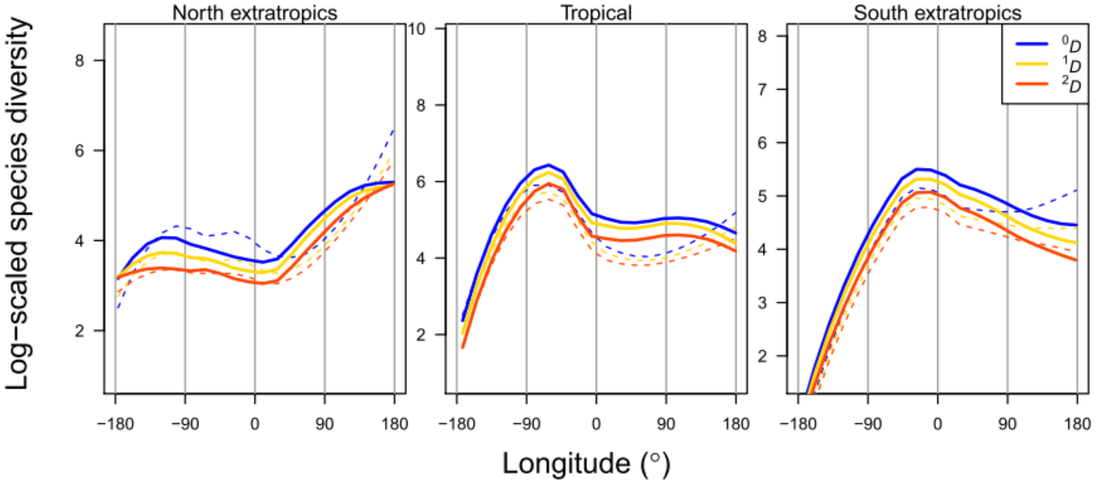


Longitudinal gradient of species diversity based on Hill numbers ($^qD$; $q$ = 0, 1, 2) at approximately 100 km × 100 km grid cell level. Dashed and solid lines represent observed and standardised (at 40th percentile of sample coverage = 0.82) values, respectively. Loess (locally estimated scatterplot smoothing) curves (scaling parameter alpha = 0.6) are shown

**Fig. S21.**

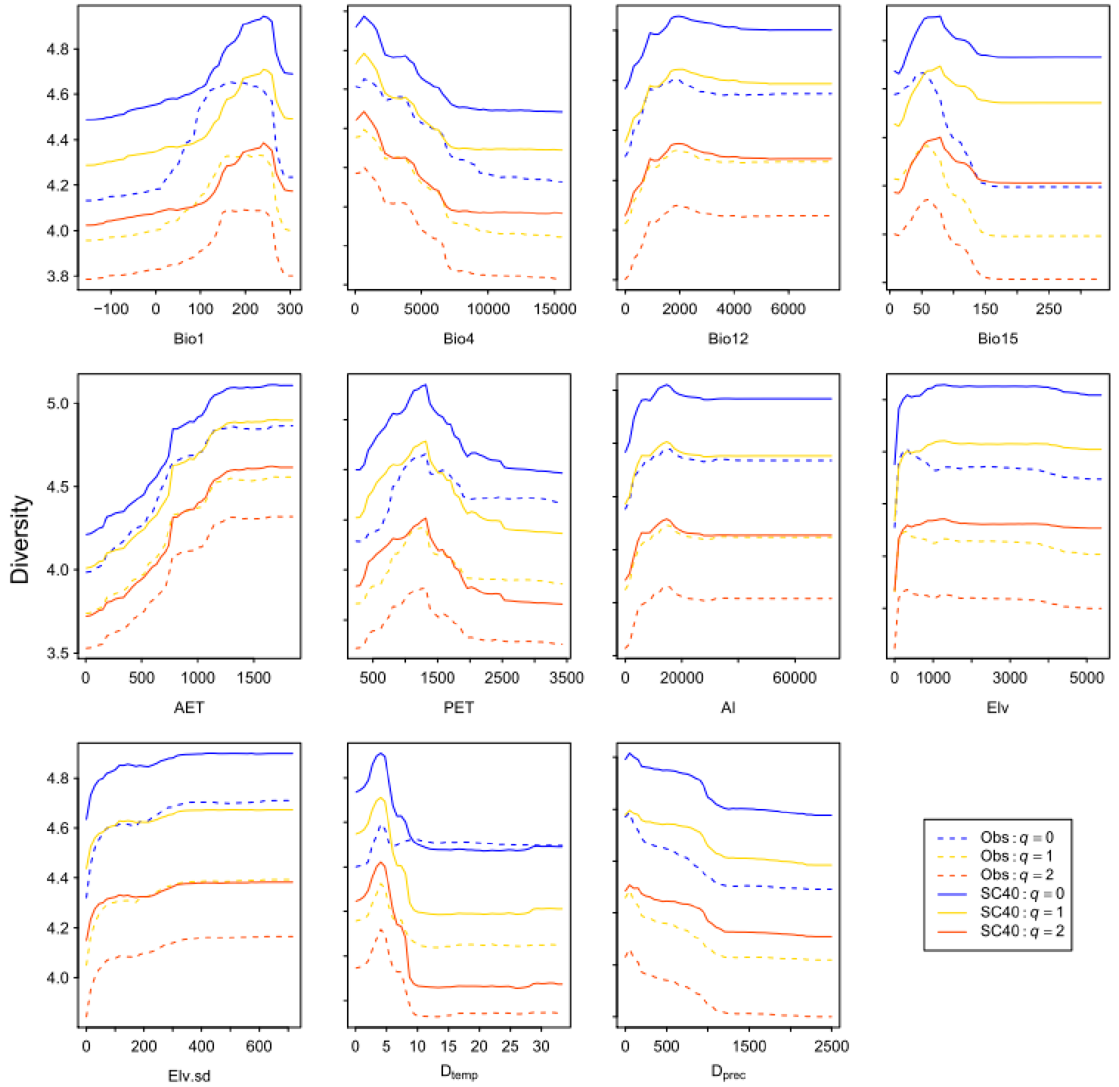


Comparison of partial dependency of explanatory variables in the Random Forest models explaining species diversity based on Hill numbers (${}^qD$; $q$ = 0, 1, 2) at approximately 100 km × 100 km grid cell level. Observed diversity (Obs) and standardized (at 40th percentile of sample coverage = 0.82) species diversity (SC40) are shown.

**Fig. S22.**

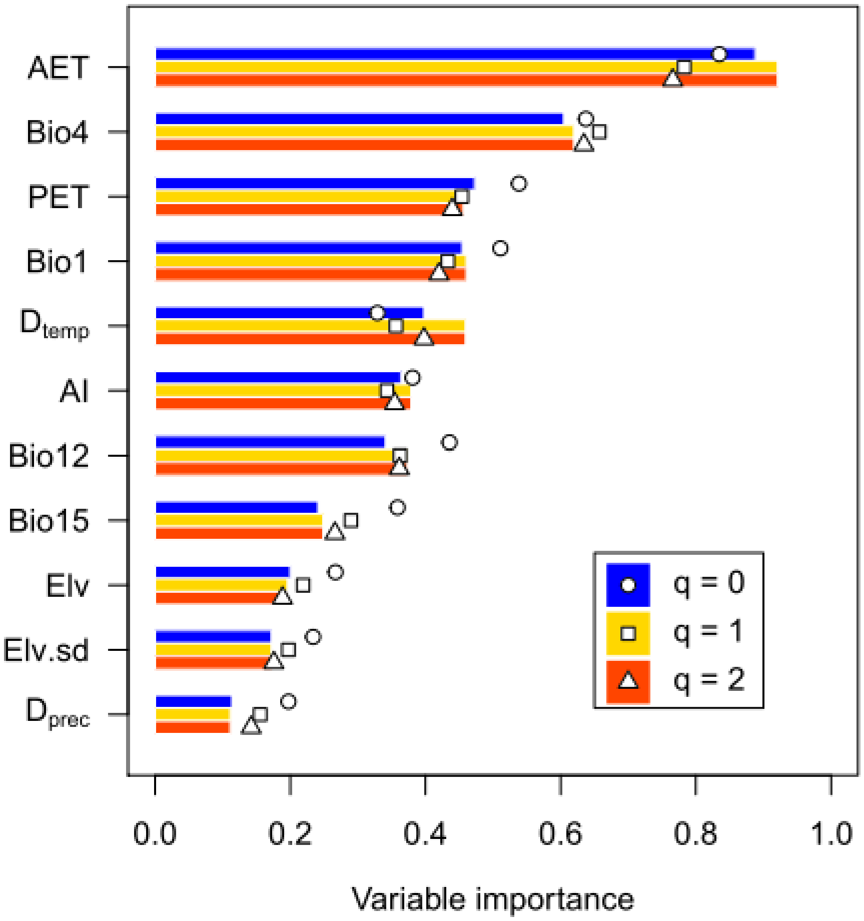


Comparison of relative importance of explanatory variables in the random forest models explaining species diversity based on Hill numbers ($^{q}D$; $q$ = 0, 1, 2) at approximately 100 km × 100 km grid cell level. Bars and points represent observed and standardised (at 40th percentile of sample coverage = 0.82) values, respectively.

**Fig. S23.**

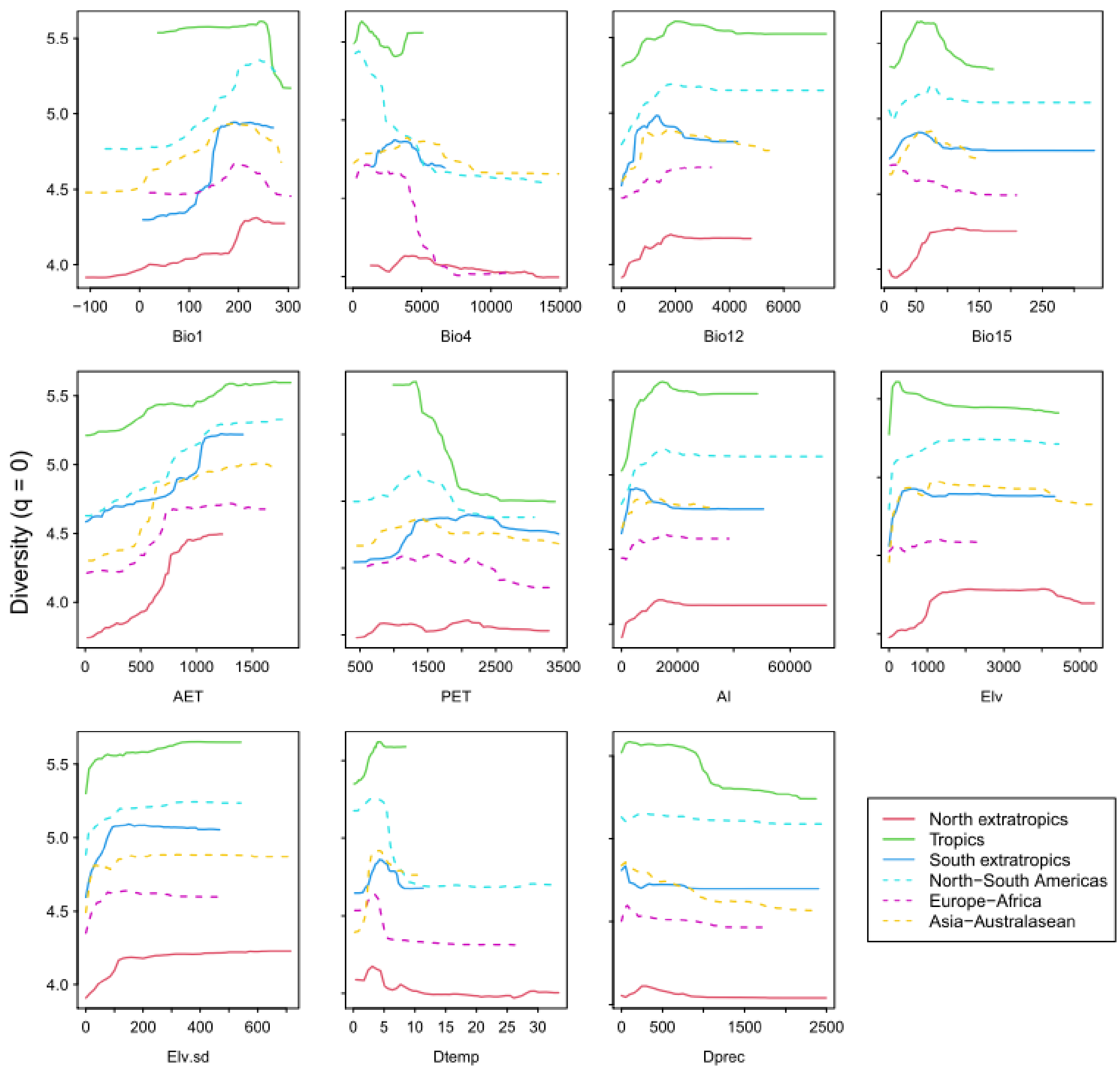


Comparison of partial dependency of explanatory variables in the random forest models explaining species richness among latitudinal (tropics and north and south extratropics) or latitudinal (Americas, Europe to Africa, and Asia to Australia) zones. The species richness at approximately 100 km × 100 km grid cell was standardised at the 40th percentile of sample coverage (0.82). The environmental variables are mean annual temperature (Bio1), temperature seasonality (Bio4), annual precipitation (Bio12), precipitation seasonality (Bio15), actual evapotranspiration (AET), potential evapotranspiration (PET), aridity index (AI), average elevation (Elv), standard deviation of elevation (Elv_sd), and differences in temperature ($D_{temp}$) and precipitation ($D_{prec}$) between the Last Glacier Maxima and the present.

**Fig. S24.**

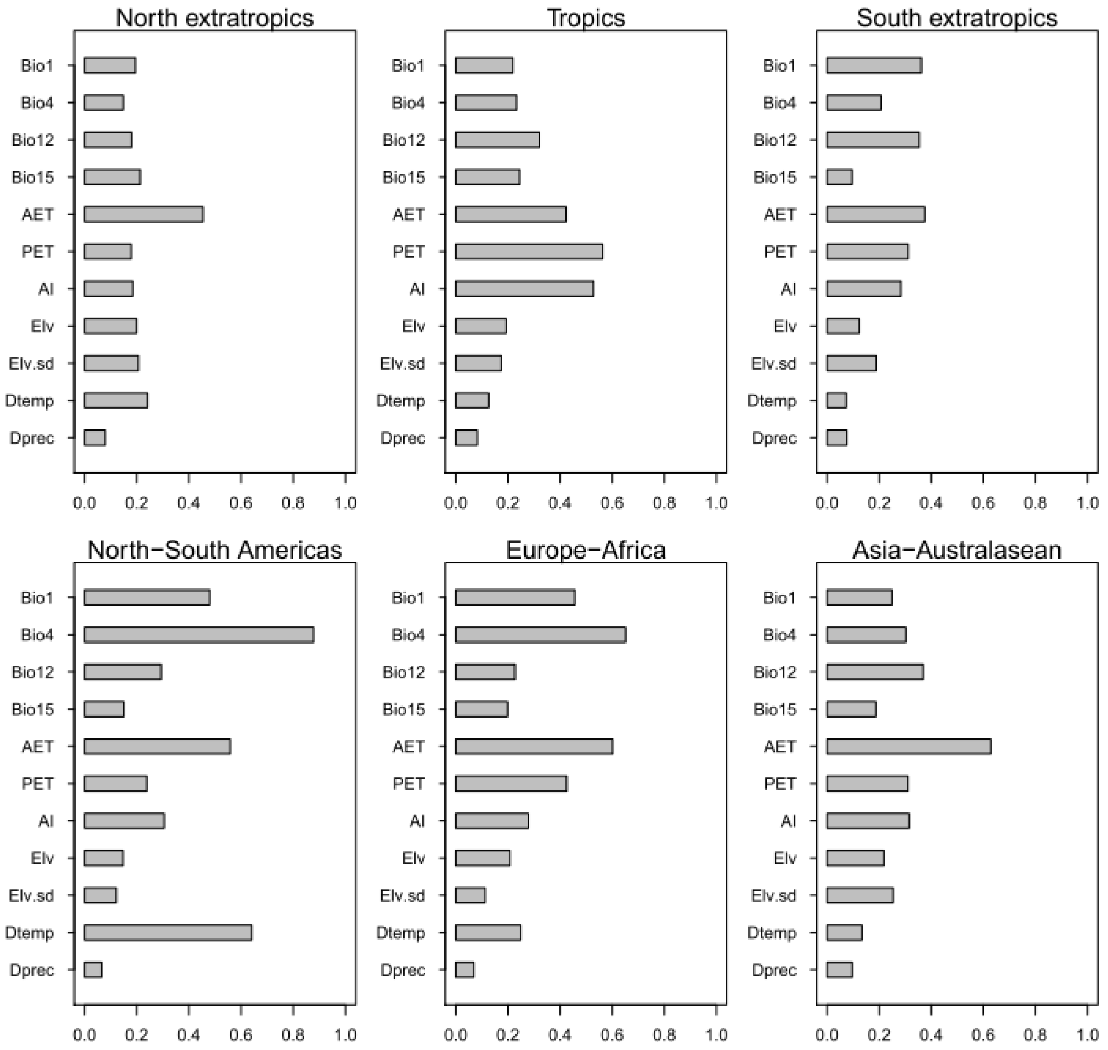


Comparison of relative importance of explanatory variables in the random forest models explaining species richness among latitudinal (tropical and north and south extratropics) or latitudinal (Americas, Europe to Africa, and Asia to Australia) zones. The species richness at approximately 100 km × 100 km grid cell was standardised at the 40th percentile of sample coverage (0.82). The environmental variables are mean annual temperature (Bio1), temperature seasonality (Bio4), annual precipitation (Bio12), precipitation seasonality (Bio15), actual evapotranspiration (AET), potential evapotranspiration (PET), aridity index (AI), average elevation (Elv), standard deviation of elevation (Elv.sd), and differences in temperature ($D_{temp}$) and precipitation ($D_{prec}$) between the Last Glacier Maxima and the present.

**Fig. S25**

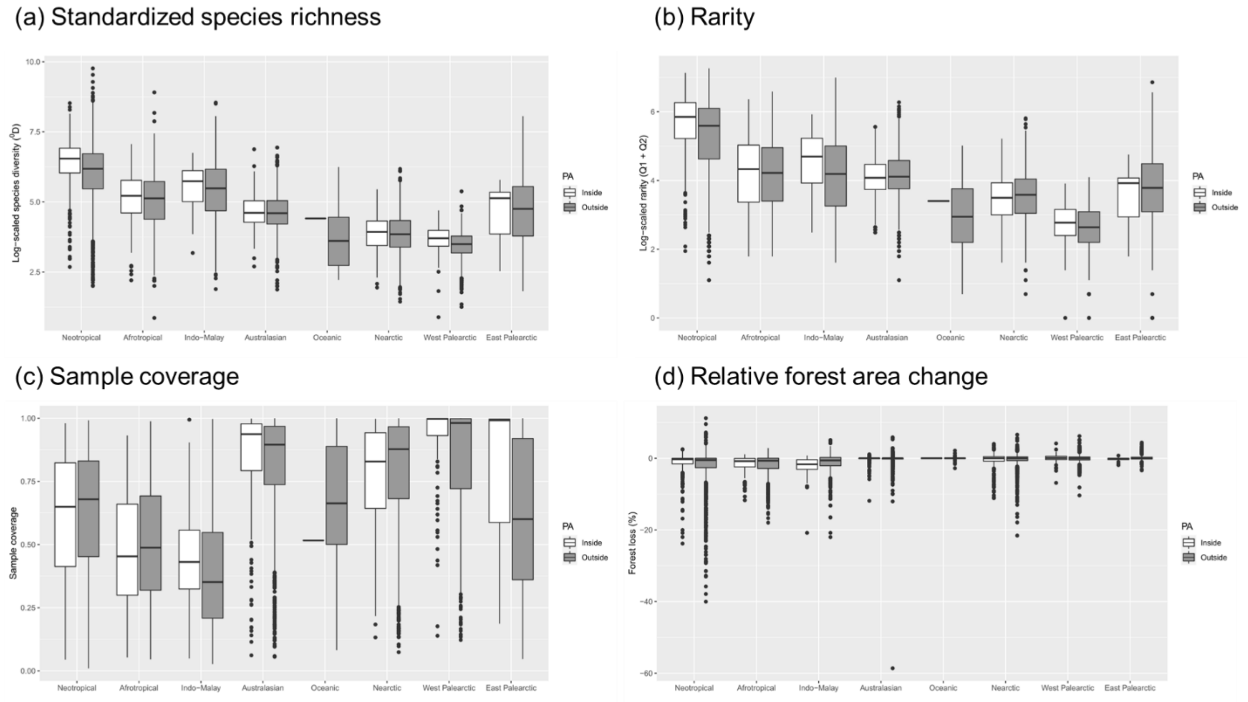


Comparison of grid-cell properties associated with woody angiosperm diversity between inside and outside the existing protected areas (PA): (a) log-scaled standardized species richness, (b) rarity defined as the number of unique and duplicated species, (c) sample coverage, and (d) relative forest area changes between 2000 and 2020.

**Fig. S26**

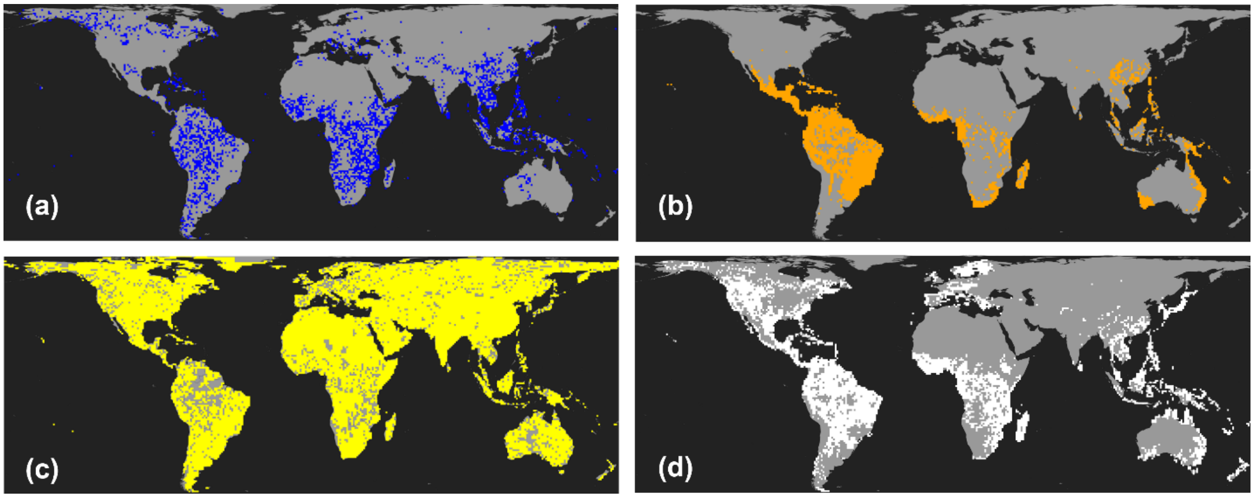


Global map of binarized variables used to determine spatial priority areas for improving sampling completeness of species occurrence records of woody angiosperms: a) low sampling coverage areas (lower than the 30th percentile; blue); high species rarity areas (upper than the 70th percentile; orange); unprotected areas (yellow); deforested areas where forest cover was lost during 2000-2020 (white).

**Table S1.**

Overview of literature sources used to compile the woody angiosperm data

| Type | Sources |
|---|---|
| Country flora list | Refs. 83-180 |
| Botanical literature | Refs. 88, 93, 99, 100, 117, 140, 145, 171, 182-206 |
| Species occurrence records | Refs. 114, 207-211; GBIF* |

* See table S3 for doi for the occurrence data downloads

**Table S2 (separate file)**
Global woody angiosperm species list

**Table S3 (separate file)**

Digital object identifiers (doi) for the download of species occurrence data from GBIF

**Table S4**

Sample coverage (SC) values at 1st, 5th, 10th, 20th, 30th, 40th and 50th percentiles at different spatial resolution

| Spatial resolution | SC1 | SC5 | SC10 | SC20 | SC30 | SC40 | SC50 |
|---|---|---|---|---|---|---|---|
| ca 100 km × 100 km | 0.142 | 0.313 | 0.431 | 0.596 | 0.729 | 0.820 | 0.889 |
| ca 200 km × 200 km | 0.227 | 0.449 | 0.579 | 0.734 | 0.836 | 0.905 | 0.944 |
| ca 400 km × 400 km | 0.336 | 0.559 | 0.697 | 0.829 | 0.902 | 0.95 | 0.972 |
| ca 800 km × 800 km | 0.470 | 0.69 | 0.799 | 0.896 | 0.951 | 0.972 | 0.986 |
